# Single-cell dynamics reveal a stress-induced decision between cardiomyocyte hypertrophy and apoptosis

**DOI:** 10.1101/2021.03.17.435783

**Authors:** Bryan Chun, Lavie Ngo, Jeffrey J. Saucerman

## Abstract

Cardiomyocyte hypertrophy and apoptosis underlie cardiomyopathies and heart failure. While previous studies have reported both hypertrophy and apoptosis at the population level, how individual cells commit to these distinct analog and digital fates is unclear. To elucidate how individual cells decide to grow and/or die, we developed a high-content microscopy approach to track single-cell cardiomyocyte dynamics. Even untreated cells exhibited substantial single-cell variability in growth and death. Uniform treatments of staurosporine or phenylephrine induced distinctive morphological programs resulting in apoptosis and hypertrophy, respectively, but only in cell subpopulations. Increasing concentrations of the β-adrenergic receptor agonist isoproterenol caused a population-level biphasic induction of hypertrophy and then apoptosis, consistent with either apoptosis in the most hypertrophic cells (a grow-and-die model) or an early decision between hypertrophy and delayed apoptosis (a grow-or-die model). By tracking single-cell fates, we found that when stressed with either isoproterenol or phenylphrine, individual cells that hypertrophy are protected from later apoptosis. Further, caspase 3 inhibition shifted the single-cell probability from apoptosis to hypertrophy fates. Machine learning models found that a cell’s initial size and DNA content or condensation could predict a cell’s bias for hypertrophy or apoptosis. Together, these data support a grow-or-die conceptual model for cardiomyocyte decisions. This single-cell profiling method for tracking joint analog-digital cell decisions reveals that despite hypertrophy and apoptosis co-occurring at the population level, individual cardiomyocytes decide early whether to grow or die.

## Introduction

Cells utilize a finite number of signaling pathways and intermediates to determine a multitude of cell fates such as developmental specification [1], [2] or drug responses [3], [4], [5]. For example, heart failure involves both cardiomyocyte hypertrophy and apoptosis [6], [7], [8]. While apoptosis has been clearly implicated in heart failure progression [9], hypertrophy can be physiological or pathological [10], [11]. Furthermore, apoptosis and hypertrophy may have interacting roles that result in the transition from physiological to pathological hypertrophy. Understanding the differential co-regulation of hypertrophy and apoptosis is critical to differentiate between pathological hypertrophy, which is accompanied by apoptosis, and compensatory hypertrophy in the absence of apoptosis [12], [13], [14], [15]. These interactions need to be better understood to develop targeted therapeutic strategies for heart failure [16].

A number of studies have shown that cardiomyocyte hypertrophy and apoptosis are co-regulated *in vivo.* For example, pressure-overload and catecholamine overstimulation induce both hypertrophy and apoptosis [17], [18], [19]. Calcineurin overexpression elicits hypertrophy that is protective against apoptosis [20], even though calcineurin directly dephosphorylates the pro-apoptotic Bcl-2 family protein BAD [21], [22]. Despite insight into population-level responses, *in vivo* studies have not elucidated the dynamic, functional relationships between hypertrophy and apoptosis at the level of single cells.

One recent technology for capturing single-cell heterogeneity is single-cell RNA sequencing, which has provided a number of important insights into health and disease [3], [4], [23]. For example, Chaffin et al. identified three distinct populations of cardiomyocytes that are differentially distributed in human hearts with dilated and hypertrophic cardiomyopathies [24].

Other studies have used single-cell RNA sequencing (scRNA-seq) to characterize cell subpopulations that regulate development, differentiation, and in response to drugs that could not be resolved with bulk measurements at the population level. Still, for most pathological responses in the heart, the dynamic and causal relationships between these co-occuring subpopulations as they respond to perturbations remains unclear [12], [25].

In this work, we propose two simple conceptual models for joint cardiomyocyte decision-making of hypertrophy and apoptosis. A “grow-and-die” model represents serial decision-making, where cells hypertrophy but then later apoptose due to an inability to handle growth stresses [26]. In contrast, the importance of caspase-mediated apoptosis but also reported sub-apoptotic caspase activity in hypertrophy [27] is consistent with a distinct parallel decision-making “grow-or-die” model based on a threshold of caspase activity [27], [28]. However, both *in vivo* and *in vitro* [29], [30], [31], [32], [33], [34] experiments have examined fixed timepoints at the population level, which does not provide the single-cell tracking that is required to discriminate between “grow-or-die” vs. “grow-and-die” conceptual models. To overcome these limitations, here, we designed a live-cell high-content microscopy approach with machine learning image analysis to track the dynamics of individual neonatal rat cardiomyocyte growth and death simultaneously. Using this high-content microscopy platform, we characterize the single-cell dynamic hypertrophy and apoptosis responses of cardiomyocytes to multiple perturbations that support the “grow-or-die” model of decision making, together with a machine learning model that identifies cell size, shape, and nuclear features that predict single-cell commitment to hypertrophy or apoptosis. Our work thus develops an experimental-machine learning framework to reveal how cells interpret stress signals based on their initial state to guide single-cell decisions of growth or death.

## Results

### Assay design for single-cell phenotyping of hypertrophy-apoptosis dynamics

We developed a live single-cell microscopy assay to track multiple simultaneous fluorescent readouts of hypertrophic and apoptotic activity (**Figure 1A**). Neonatal rat cardiomyocytes were transduced with an eGFP-expressing lentivirus as a live-cell reporter of single-cell morphology and protein synthesis. Nuclei were labeled with Hoechst, and general cell death and apoptosis were measured with propidium iodide (PI) and fluorescently-conjugated annexin V (A5). To measure live-cell dynamics, individual wells in a 96-well microplate representing different treatment conditions were imaged for 48 hours with a 1 hour resolution (**Figure 1B**). After live-cell imaging, cells were fixed, permeabilized, and immunostained to detect cardiomyocyte-specific ɑ-actinin expression.

**Figure 1.**
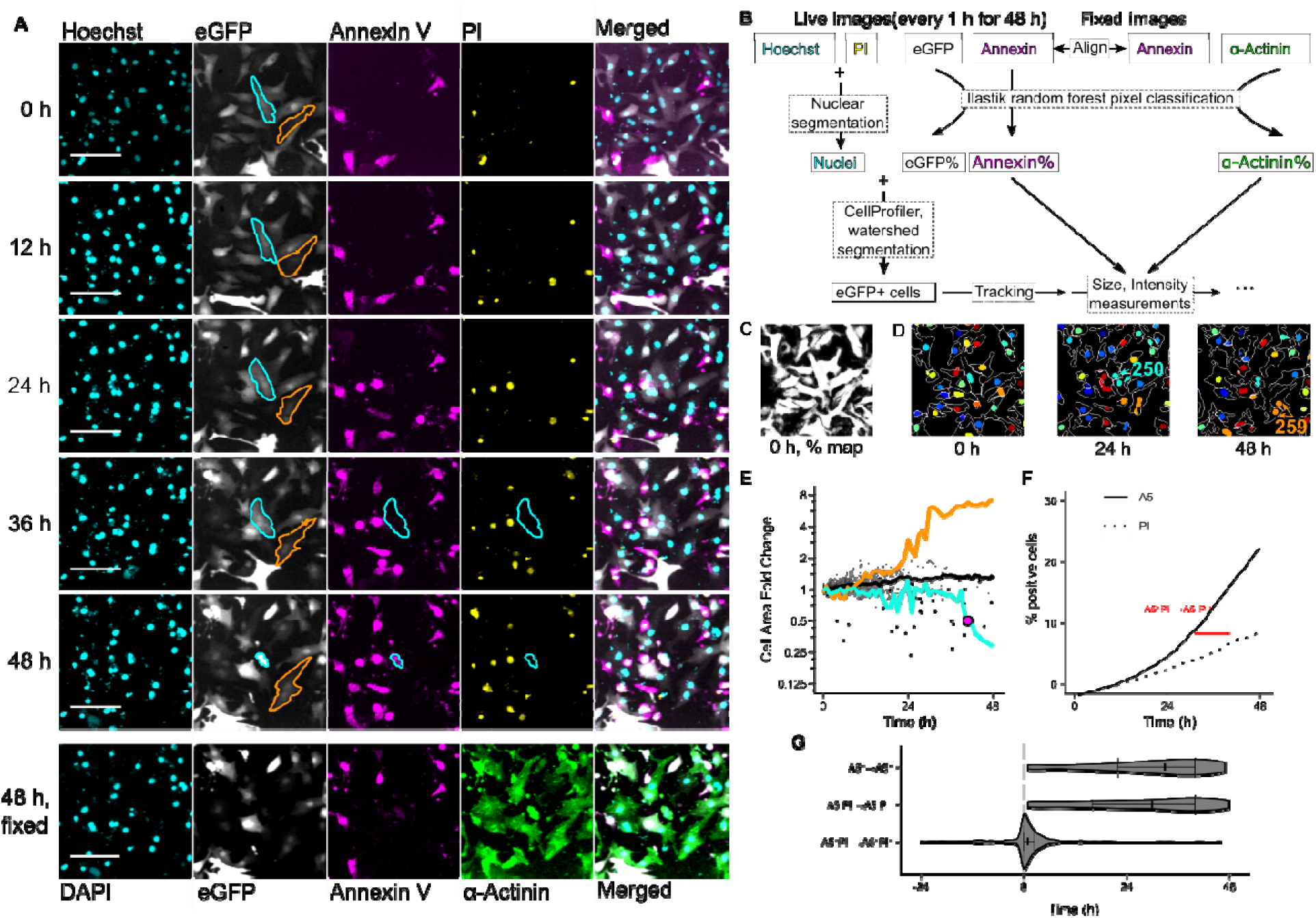
High-content time-lapse microscopy of single-cell morphological and apoptosis dynamics of neonatal rat cardiac myocytes. (**A**) Representative timelapse images of neonatal rat cardiomyocytes treated with 0.1% DMSO (neg. control). Live imaging tracked changes in morphology of eGFP-expressing cells and apoptotic A5 and PI binding over 48 hours (rows 1 through 5). After two days (row 6), cells were fixed and stained for DNA (DAPI, cyan) and α-actinin (green). Outlines represent manual segmentation of cells undergoing hypertrophy (orange) or early apoptotic A5 binding and PI exclusion (cyan). Scale bar is 100 μm. (**B**) Schematic of image analysis pipeline for single-cell cardiomyocyte hypertrophy and apoptosis dynamics. (**C**) Example eGFP fluorescence probability map for the GFP image in (A) for image thresholding and segmentation. (**D**) Example segmentation and tracking of cells in (A). White outlines represent automated segmentation results of eGFP cells and pseudo-coloured ovals represent nuclear segmentation. The tracking of the two outlined cells in (A) are marked with their identifier (cyan and orange numbers and arrows). (**E**) Single-cell heterogeneity in cardiomyocyte hypertrophy dynamics (black lines). Orange and cyan traces correspond to previously identified cells. Solid black line corresponds to the population median. Black dots indicate onset of A5 binding as does the large magenta dot for the cyan trace. Cell Area Fold Change reported as cell areas normalized to the average of areas measured at initial two timepoints. (**F**) Empirical cumulative distribution function of A5 (solid line) and PI (dashed line) binding for all DMSO-treated cells (n > 3600 cells from 3 wells from each of 2 independent isolations). The rightward shift of the PI binding curve indicates later onset of PI binding in the cell population consistent with apoptosis. (**G**). Single-cell quantification of onset of A5 and PI binding show that PI occurs after A5 binding, with the lag between those two events quantified (red distribution; * p < .0001, one-sample Wilcoxon signed rank test, n > 450 cells).

We developed machine learning image analyses to measure dynamics of multiple metrics related to cellular morphology, viability, and final ɑ-actinin expression at a single-cell level (**Methods**). To correct for cell-to-cell eGFP expression variability and uneven fluorescence that persisted after flatfield illumination, we trained a random forest classifier to generate probability maps of eGFP (**Figure 1C**). The random forest model was robust to at least a 5-fold range in variations in fluorescence signal-to-noise ratio, allowing optimization of fluorescence acquisition settings to minimize phototoxicity (**Supplemental Figure 1**). We used this eGFP probability map for thresholding, single-cell segmentation, and tracking for longitudinal morphology and fluorescence measurements (**Figure 1D** and **Methods**). As our eGFP reporter uses a ubiquitous CMV promoter that does not differentiate between cardiomyocytes and other cell types, we tracked cardiomyocytes by first live-cell tracking in eGFP and then colocalizing eGFP with the α-actinin probability map in the fixed images. Similarly, a live A5 probability map was used to determine A5 positivity of each cell.

Our assay was first validated with cells treated with a DMSO negative vehicle control. Even in the absence of hypertrophic or apoptotic stimuli, there was considerable single-cell variability of both hypertrophic (**Figure 1A** and **D**, orange outline and traces) and apoptotic trajectories (cyan outline and traces). Live single-cell tracking of the DMSO condition across 48 hours is shown in **Supplemental Movie 1**. At the cell-population level, A5 binding accumulated earlier than PI uptake, indicating that most of the death observed was due to apoptosis rather than other death processes (**Figure 1F**). This apoptotic lag between initial A5 binding and later PI uptake at the population level was validated in individual A5/PI double-positive cells (**Figure 1G**).

### Cellular and nuclear profiling of staurosporine-induced apoptosis dynamics

Apoptosis is characterized by morphological and biochemical changes to both cell and nucleus. As a representative cellular perturbation to examine apoptosis dynamics with our high-content imaging assay, we used the broad spectrum ATP-competitive kinase inhibitor staurosporine (STS), which is well-known to induce apoptosis in cardiomyocytes and other cell types [35], [36], [37], [38]. STS-induced apoptotic or death markers such as A5 or PI have been characterized in fixed cardiomyocytes at a coarse time resolution [38], [39]. Our high-content imaging assay uniquely allows for simultaneous characterization and high-resolution dynamic single-cell tracking of multiple features related to apoptosis including A5 binding, PI uptake, changes in nuclei morphology, and chromatin condensation and degradation.

Treatment with STS induced substantial A5 binding and PI uptake compared to DMSO vehicle-control treatment (**Figure 2A** and **B**). Consistent with prior studies, bulk-population dynamics of total accumulation of A5 and PI were again indicative of apoptosis, with PI-uptake occurring later than that of A5 (**Figure 2C**). However, single-cell dynamics of apoptosis were markedly different between STS and DMSO treatments. DMSO treatment induced a biphasic response with initially low levels of A5 and PI binding between 0 and 24 hours and a second phase of increased apoptotic activity after 24 hours. In contrast, STS-treated cells largely completed apoptosis within the first 36 hours of treatment, as shown in **Supplemental Movie 2.** Single-cell tracking revealed a time lag from A5+/PI- to A5+/PI+ (**Figure 2D**), consistent with apoptosis and not other forms of cell death. This indicates the added value of measuring single-cell dynamics of cell death.

**Figure 2.**
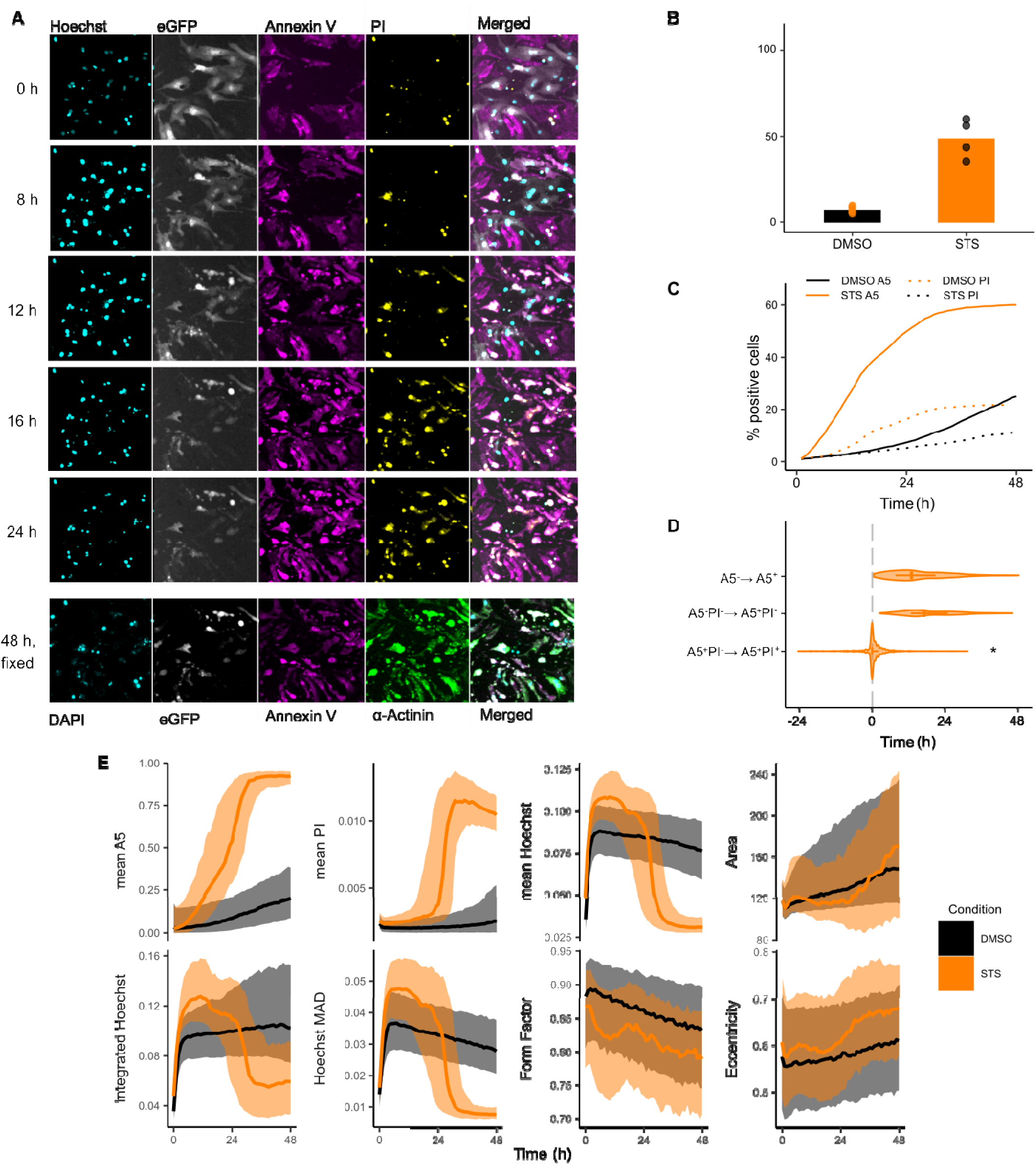
Single-cell dynamics show that cardiomyocyte death induced by staurosporine follows a sequence of nuclear permeability and shape, chromatin condensation, annexin 5 binding, and membrane permeability. (**A**) Representative timelapse images of neonatal rat cardiomyocytes treated with 1 μM STS show induction of A5 and PI binding (magenta and yellow, respectively) and pyknosis (cyan). (**B**) 1 μM STS increased the fraction of apoptosis (A5+/PI-) cardiomyocytes compared to the 0.1% DMSO vehicle control condition. ** p = 0.0049, Welch’s unpaired *t*-test, 2-3 wells per condition for each of n = 2 independent cell isolations (wells represented as jitter points). (**C**) Empirical cumulative distribution function of A5 (solid) and PI (dashed) binding for all STS-treated cells. Same DMSO traces as in Figure 1F. (**D**) Single-cell quantification of time lag from A5+/PI- to A5+/PI+ in response to STS treatment (* p < 10^-11^, one-sample Wilcoxon signed rank test, n >1100 cells). (**E**) Single-cell dynamics of nuclear metrics in response to either 1 μM STS or 0.1% DMSO control. Solid lines indicate population median, shaded areas indicate interquartile range.

Next, we sought to characterize the temporal dynamics of additional cellular features that may be associated with apoptosis (**Figure 2E**). As expected, STS treatment induced global increases of cell membrane A5 binding followed by nuclear PI uptake. Motivated by known apoptotic nuclear and chromatin changes [40], we also investigated dynamic changes in nuclear Hoechst staining. Mean and total (integrated) Hoechst intensity (reflecting membrane permeability), and median absolute deviation (MAD) of the Hoechst intensity (reflecting chromatin condensation [40]) all exhibited coordinated increases with STS-treatment [40], [41]. After approximately 24 hours, STS-treated trajectories of Hoechst decreased below that of DMSO-treated trajectories, consistent with DNA degradative apoptotic processes. We also investigated changes in nuclear morphology. During the first 36 hours, nuclei of STS-treated cells had smaller nuclei, which later increased in size relative to their DMSO-treated counterparts. Interestingly, nuclear form factor decreased and eccentricity increased in STS-treated cells relative to DMSO-treated cells even at the very early times, indicating more irregular and less circular nuclei consistent with apoptosis. Overall, single-cell tracking of multiple cellular features delineates a temporal sequence of apoptotic kinetics, starting with changes to nuclear permeability and shape, followed by A5 binding, and finally loss of cell membrane integrity.

### Cellular profiling of phenylephrine-induced hypertrophy dynamics

To examine hypertrophy dynamics with our high-content imaging assay, we stimulated cardiomyocytes with the catecholamine phenylephrine (PE), an ɑ1-adrenergic agonist and a well-known inducer of cardiomyocyte hypertrophy [42], [43]. Cardiac ɑ1-adrenergic signaling has also been shown to activate cell survival signaling pathways [32], [44]. We further examined the protein translation inhibitor cycloheximide (CHX) as an additional negative control to control for baseline protein synthesis [45].

Treatment with PE increased two established metrics of hypertrophy: an increase in cell area and hypertrophy and eGFP expression (**Figure 3A**-**C**). We previously found PE-induced increase in cardiomyocyte 2D cell area to strongly correlate with 3D cell volume [43], and eGFP expression driven by a constitutive promoter reflects overall protein synthesis. The greatest increase in median cell areas relative to DMSO occurred around 24 h (**Figure 3D**). Interestingly, there was substantial single-cell variability in PE-induced changes in cell area. Comparison of the empirical cumulative distribution functions (eCDFs) of the DMSO- and PE-treated cells show not simply a rightward shift in distribution when cells are treated with a hypertrophic agonist, but rather an increase in the upper tail of distribution of cell areas (**Figure 3F**). This indicates that population-level increases in cell area seen with PE treatment are due to a subpopulation of hypertrophic responders as opposed to global increases in cell area. PE treatment increased the proportion of hypertrophic responders, or cells with fold change areas greater than that of the 75th percentile of DMSO, while CHX treatment reduced this proportion (**Figure 3G**). Cells treated with PE are shown in **Supplemental Movie 3** with increases in cell area and GFP expression on the single-cell level.

**Figure 3.**
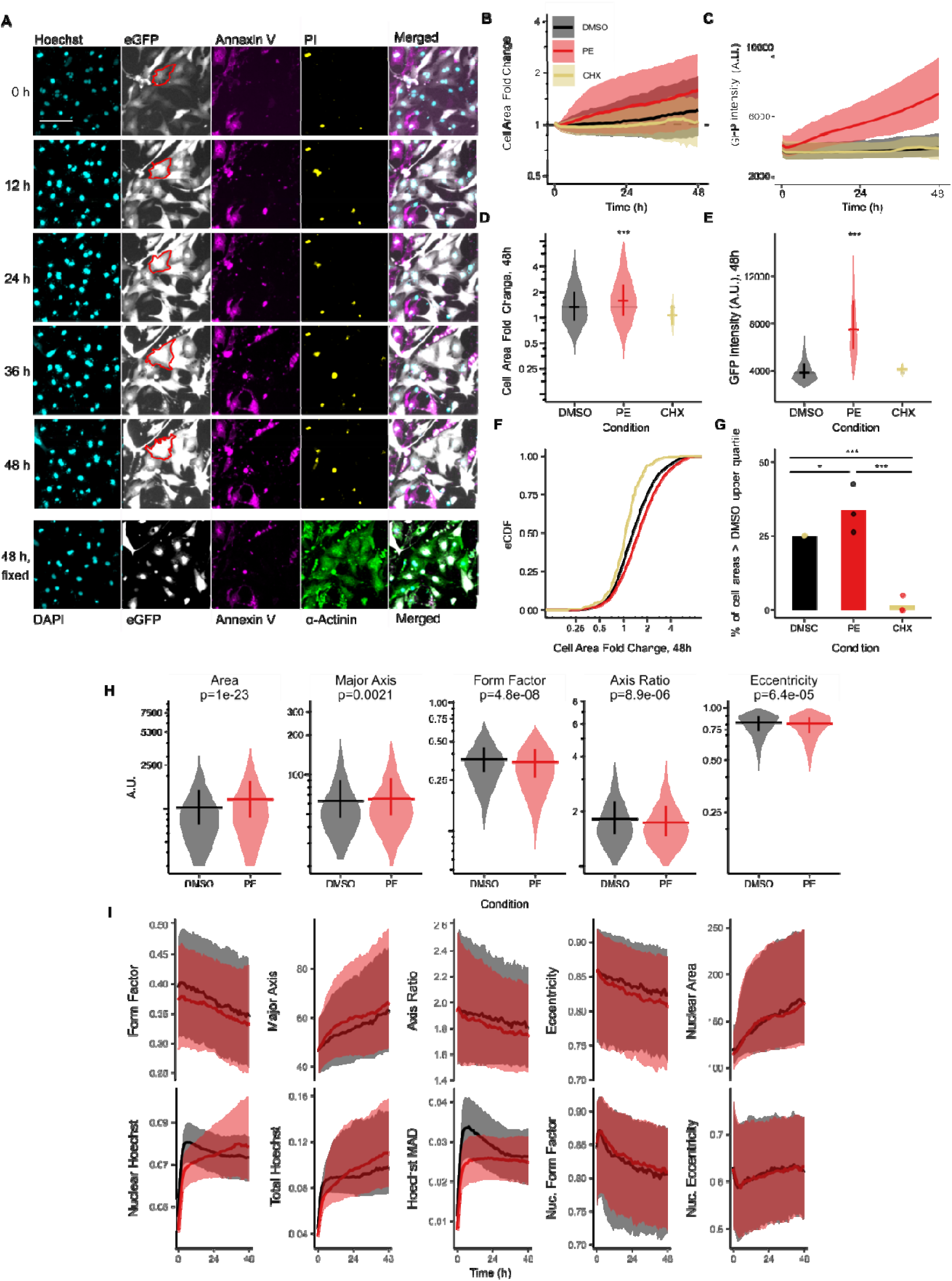
Phenylephrine induces coordinated changes in cell morphology, protein synthesis, and nuclear integrity. (**A**) Representative timelapse images of neonatal rat cardiomyocytes treated with 50 μM PE showing induction of hypertrophy (red outline) and eGFP expression (gray). 48 h hypertrophy dynamics (**B**) and eGFP dynamics (**C**) of cells treated with 50 μM PE, 50 μM CHX, or 0.1% DMSO vehicle control. Solid lines indicate population median, shaded areas indicate interquartile range. 48 h treatment with 50μM PE induces hypertrophy (**D**) or eGFP expression (**E**). Errorbars indicate median and interquartile range *** p < 2*10^-16 *vs.* DMSO control, Kruskal–Wallis H test followed by one-sided Mann–Whitney U post-hoc test with Benjamini-Hochberg multiple testing correction, n >2000 cells. (**F**) Empirical cumulative distribution functions (eCDFs) of cell areas. Same data as in (**D**). (**G**) Percent of cells with areas greater than DMSO 75th percentile. Same data as in (**D**). *** p ≤ 1.58*10^-6, mixed-effects ANOVA followed by Tukey HSD, 6 wells per condition for each of n = 3 independent cell isolations (represented as jitter points). (**H**) Cell morphology metrics of cells treated with either 50 μM PE or 0.1% DMSO vehicle control at 48 h. Solid lines indicate population median, and error bars indicate interquartile range. p-values represent Kruskal–Wallis H test followed by one-sided Mann–Whitney U post-hoc test with Benjamini-Hochberg multiple testing correction, n > 2000 cells. (**I**) Single-cell dynamics of cellular or nuclear metrics in response to either 50 μM PE or 0.1% DMSO control. Solid lines indicate population median, shaded areas indicate interquartile range.

We previously identified divergent characteristics of cellular hypertrophy induced by different hypertrophic agonists [46]. While our previous approach segmented cells based on ɑ-actinin immunofluorescence, here we found greater fidelity when further incorporating cell-to-cell variation in eGFP expression (**Supplemental Figure 2**). We therefore re-characterized the same metrics of cell shape and size and found that treatment with PE led to significant changes to all metrics observed (**Figure 3H**). As expected, PE increased raw cell area and major axis length. Elongation (axis ratio), eccentricity, and form factor all decreased, indicating PE-treated cells became more circular but more spiky (**Figure 3A**, red outline).

Previous reports have suggested a role for increased DNA synthesis and polyploidy in cardiac hypertrophy, and we recently published a method for determining cardiomyocyte nuclear ploidy with the nuclear stain DAPI [47], [48]. Further, Hoechst has been shown to be a reliable marker for chromatin condensation, which may play a functional role in transcription [49]. We therefore asked whether the hypertrophic stimulus PE also affected nuclear characteristics as measured by Hoechst. While nuclear morphological characteristics remained unchanged, Hoechst MAD was decreased, indicating altered chromatin condensation. We found that PE-treated cells initially took up less Hoechst than DMSO-treated counterparts but finished with an overall brighter Hoechst signal, indicating increased DNA content (**Figure 3I**). The smaller initial Hoechst intensity may reflect more robust nuclear membrane integrity due to the cell survival signaling pathways activated by PE and the relative rates of phospholipid turnover and DNA synthesis [50], [51]. Overall, these results show that PE induces both cardiomyocyte size but also metrics indicative of nuclear integrity.

### Isoproterenol induces biphasic dose-dependent hypertrophy and apoptosis dynamics

Computational models of the cardiomyocyte hypertrophy signaling network have identified ꞵA signaling as potentially both hypertrophic and apoptotic [8], [25]. The catecholamine isoproterenol (ISO) is a non-specific β-adrenergic receptor agonist and has been separately characterized as being either hypertrophic [30], [46], [52] or apoptotic [33], [53], [54]. Our assay provides a unique opportunity to investigate the dynamics of single-cell decision-making between these co-regulated hypertrophy and apoptosis [12], [27] that cannot be resolved by snapshots at individual timepoints.

To our knowledge, our assay shows for the first time the temporal dynamics of hypertrophy and apoptosis in the same cell s, in this case with ISO treatment (**Figure 4A**). ISO was previously shown to regulate cardiomyocyte contractility and hypertrophy with distinct concentration dependence [52]. Therefore, we examined ISO responses over a 0.1-100 μM range previously examined in studies of either cardiomyocyte hypertrophy or apoptosis [32], [33], [52]. Interestingly, ISO induced a biphasic concentration-dependent hypertrophy response (**Figure 4B** and **C**). While all ISO-treated conditions induced some level of hypertrophy relative to the DMSO vehicle-treated control within 24 hours of treatment, medium to high doses of isoproterenol (10 to 100 µM ISO) induced less hypertrophy or plateaued compared to lower doses. This plateau was especially striking after 24 hours (**Figure 4D**), the timepoint associated with apoptotic responses for STS stimuli. In contrast to the biphasic hypertrophy response, A5 binding showed an ultrasensitive response with enhanced apoptosis at very high ISO concentrations (**Figure 4A**, **E**, and **F**). These data are consistent with prior population-level data from neonatal and adult rat cardiomyocytes at 10 to 100 μM ISO, which induce apoptosis as measured by cleavage of caspase 3 and PARP, TUNEL, DNA fragmentation, and regulation of ICER/Bcl2/Bad [33], [53], [55], [56], [57]. Further, such studies have shown apoptosis induced by 10 to 100 μM ISO is prevented by β-blockers and inhibitors of downstream protein kinase A, and L-type Ca^2+^ channels [33], [34], [53], [58]. Furthermore, those experiments were generally conducted in serum-free cell growth medium, which potentiate cells to undergo apoptosis [59], [60]. Consistent with previous studies on fractional killing of cancer cell lines [28], we found that 100 µM ISO induced substantial single-cell variability in kinetics of growth, apoptosis (A5 binding of PI- cells), and membrane permeability (apoptotic cells that then become PI+) (**Figure 4G**). As shown in **Figure 4G** and **H**, most apoptotic cells were smaller at the time of A5+/PI- binding (denoted as “diers”) than their predominantly hypertrophic and A5-/PI- counterparts (“survivors”). Interestingly, we found that upon apoptosis, many exhibit a rapid drop in GFP intensity but not cell area, reflecting membrane permeability **(Supplemental Figure 3).** Compared to the rates of apoptosis, there was less necrosis (A5-/PI+) in response to ISO (**Supplemental Figure 4**). This single-cell variability to a uniform stimulus indicates an unappreciated relationship between growth and death phenotypes.

**Figure 4.**
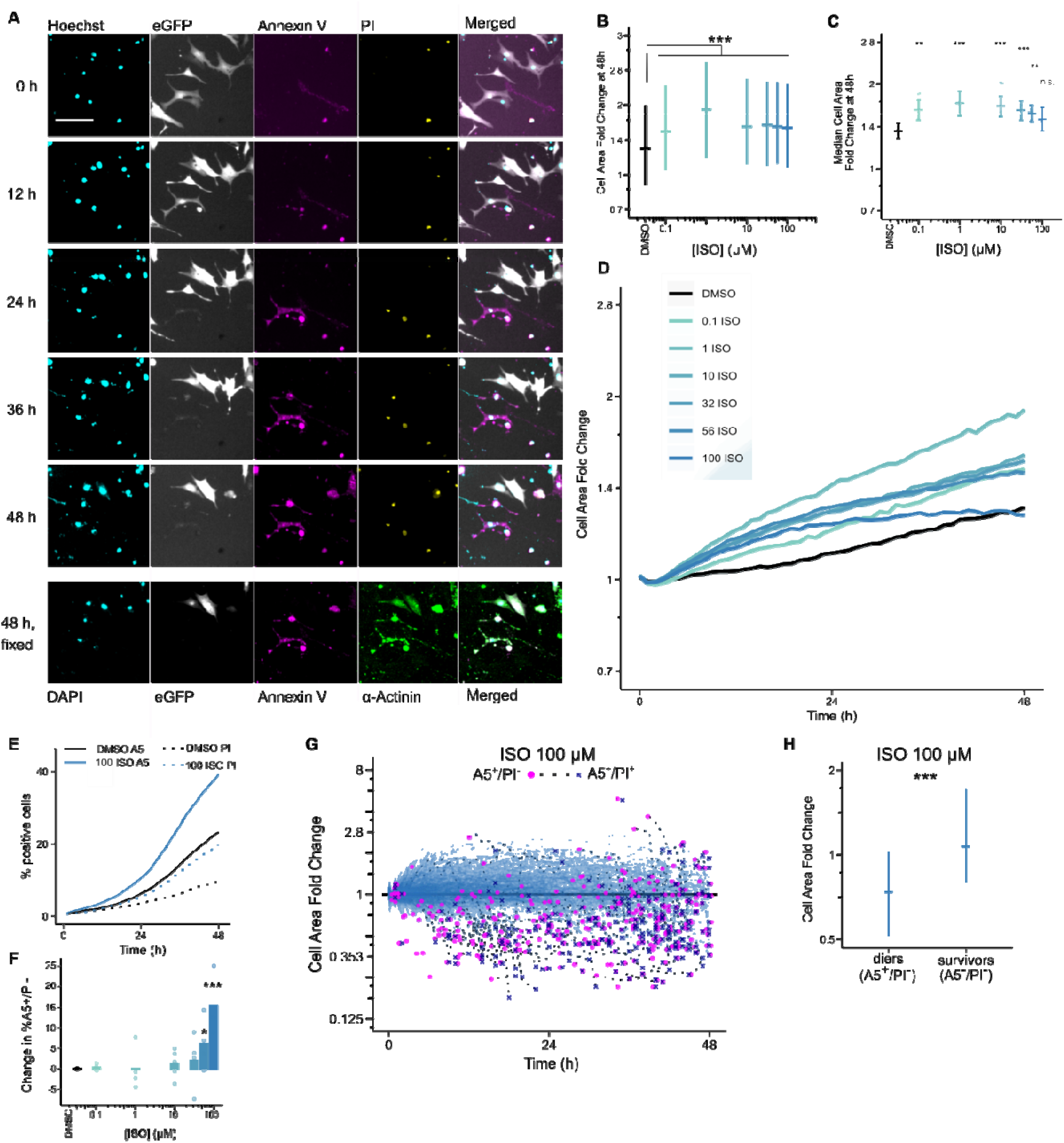
Isoproterenol stimulates distinct dose-dependent responses of hypertrophy and apoptosis, with substantial single-cell variability. (**A**) Representative timelapse images of neonatal rat cardiomyocytes treated with 100 μM ISO showing induction of hypertrophy and A5 binding (magenta). (**B**) Single-cell ISO concentration hypertrophy responses at 48 h. Solid lines indicate population median, errorbars indicate interquartile range. *** all ISO-treated conditions were significant with p ≤ 2.22*10^-16^ *vs.* DMSO control; Kruskal–Wallis H test followed by one-sided Mann–Whitney U post-hoc test with Benjamini-Hochberg multiple testing correction, n > 4000cells. (**C**) Median ISO concentration hypertrophy responses at 48 h across experiments. Error bars represent standard error of the mean. * p < 0.05, ** p < 0.01, *** p < 0.001; mixed effects ANOVA followed by one-sided Dunnett post hoc test, 6-9 wells per condition from each of n = 3-5 independent cell isolations. (**D**) ISO concentration response of hypertrophy dynamics. Shaded area represents interquartile range. **(E)** Empirical cumulative distribution function of A5 (solid) and PI (dashed) binding for all cells treated either with DMSO or 100 μM ISO. (**F**) ISO concentration response of apoposis (A5+/PI-) through 48 h. * p < .05, *** p < .001 vs DMSO control; mixed effects ANOVA followed by Dunnett post hoc test, same wells as in (**C**) with n = 3-5 independent cell isolations. (**G**) Representative single-cardiomyocyte dynamic traces of morphology of cells treated with 100 μM ISO. Magenta dots indicate onset of A5+/PI- binding, with the blue ‘X’ marker indicating any subsequent PI binding in that same cell. (**H**) Hypertrophy cell area fold change induced by 100 μM ISO at time of apoptosis (A5+/PI-,diers) or end of time course (A5-/PI-, survivors). Solid lines indicate population median, and error bars indicate interquartile range. *** p < 2.1*10^-193 *vs.* DMSO control, one-sided Mann-Whitney U test, n > 2600 cells.

### Differential response of cardiomyocyte subpopulations contributes to biphasic hypertrophy and ultrasensitive apoptosis

While hypertrophy and apoptosis co-occur in our population-level data and in stressed hearts in vivo, they logically cannot coincide within a single cell at a single time. Therefore, we proposed two distinct conceptual models of single-cell heterogeneous decision making that may reconcile this paradox. We hypothesized that single-cell decisions may be made via either 1) a “grow-and-die” model where hypertrophic stress causes apoptosis, or 2) a “grow-or-die” model, where either the local environment or initial cell state dictates cell fate (**Figure 5A**). Naturally combinations of these hypotheses or alternate hypotheses are also possible.

**Figure 5.**
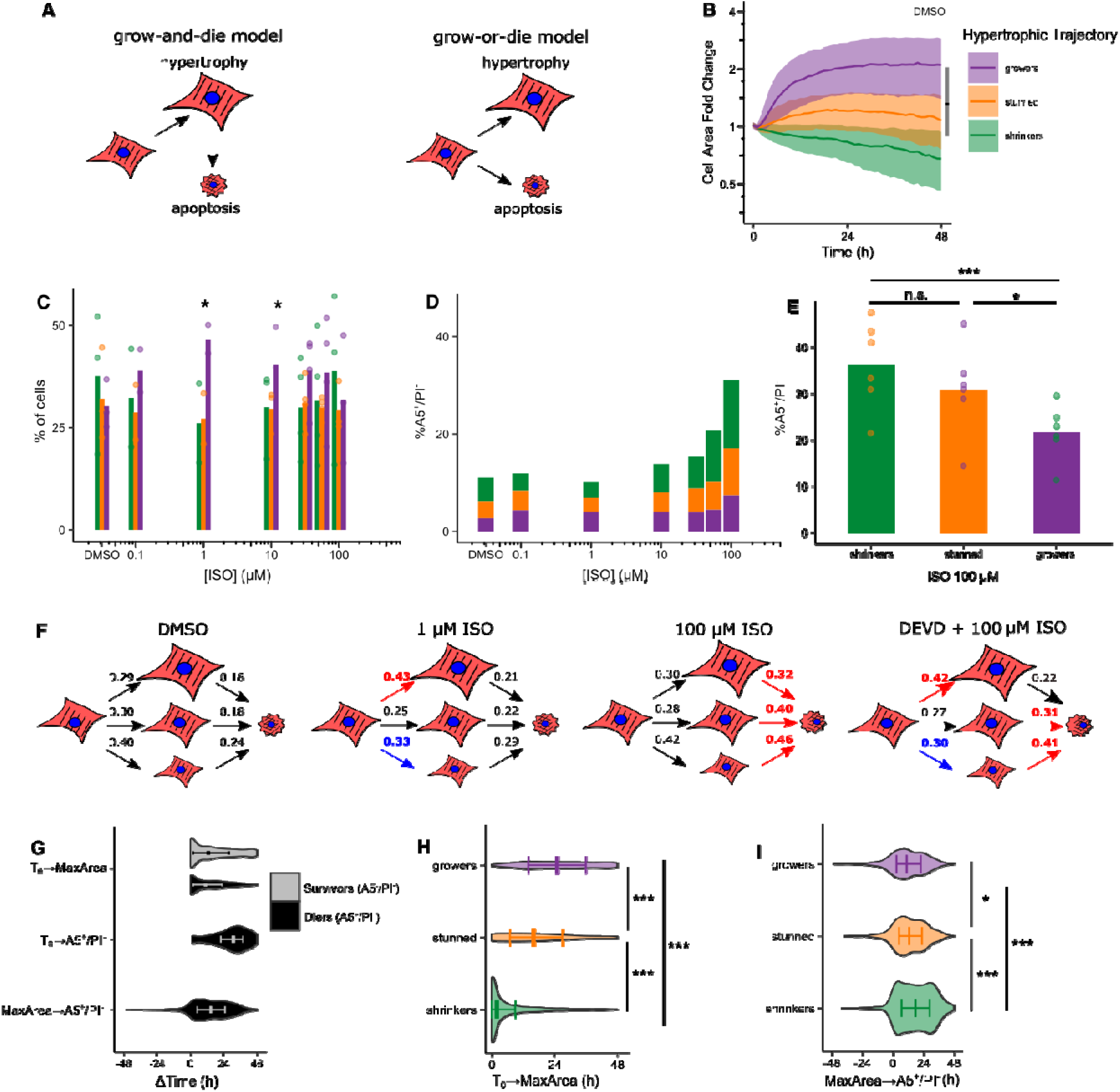
Single-cell hypertrophy-apoptosis tracking in response to isoproterenol supports a “grow-or-die” conceptual model. All data is from experiments reported in Figure 4. (**A**) Schematic of possible cardiomyocyte phenotypic decision-making models and relationships between hypertrophy and apoptosis. (**B**) The population of cells treated with 100 μM ISO was partitioned into 3 hypertrophic trajectory groups relative to the DMSO-treated population at 48 h. Growers (purple) were defined as cells whose max fold change area was greater than that of the 75th percentile of the DMSO-treated population. Stunned (orange) cells were defined as cells whose max fold change area was in the DMSO-treated interquartile range. Shrinkers (green) were defined as cells whose max fold change area was less than the 25th percentile of the DMSO-treated cells. Solid lines indicate population median, shaded areas indicate interquartile range, black line indicates population median. (**C**) 48-hour ISO concentration response according to hypertrophic trajectory. Comparisons were made between each treatment’s percent “grower” subpopulation. While the overall mixed-effects ANOVA was not statistically significant (p = 0.1399) the more powered one-sided Dunnett’s test showed statistically significant differences indicated with * p < 0.05 vs. DMSO control, 1-3 wells for each subpopulation per condition from n = 3-4 independent cell isolations. Moreover, the mixed-effects ANOVA between the 3 hypertrophy subpopulations was not significant for the DMSO treatment. There was not a significant increase in the proportion of the grower subpopulation relative to the other two subpopulations with p < 0.01, mixed-effects ANOVA followed by Tukey HSD. (**D**) Concentration response of ISO-induced apoptosis (A5+/PI-) from Figure 4E, according to hypertrophy trajectory. (**E**) Cell population makeup for 100 μM ISO. Error bars represent mean ± SEM. * p = 0.0339, *** p < 0.001, mixed-effects ANOVA followed by Tukey HSD, 1-3 wells per subpopulation from each of n = 3 independent cell isolations. (**F**) Hypertrophy and apoptosis transition state diagrams for different ISO concentrations and caspase 3 inhibitor DEVD, derived directly from panel D or **Supplemental** Figure 5 from the proportion of cells in each hypertrophic trajectory. Red or blue arrows indicate an increase or decrease in transition probability of > 0.05 compared to DMSO, respectively. (**G**) Single-cell quantification for 100 μM ISO, of onset of maximal hypertrophic area and A5 binding, and with the lag between those two events for cells that become A5+/PI- (diers) or remain A5-/PI- (survivors). For each hypertrophy subpopulation, we quantified time to maximal hypertrophy (**H**, n > 2600 cells) or time between maximal hypertrophy and A5+/PI- binding (**I,** n > 2600-4349 cells) as in **G**. Solid lines indicate population median, error bars indicate interquartile range. * p = 0.037, *** p < 2*10^-16, Kruskal–Wallis H test followed by Dunn’s All-Pairs Rank Comparison post-hoc test with Benjamini-Hochberg multiple testing correction.

To facilitate the comparison of these distinct conceptual models of hypertrophy-apoptosis decision making, we first classified cells as “growers”, “stunned”, or “shrinkers”. Strikingly, the hypertrophic dynamics of these three subpopulations identified by single-cell tracking (**Figure 5B**) are qualitatively distinct from the transient population-level hypertrophic response to 100 µM ISO (**Figure 4B**). Thus, the transient population-level hypertrophy results from the early increases in the cell area of the “growers,” followed by the later decreased cell area of the “shrinkers,” and some cells remaining “stunned”. The “shrinker” subpopulation was more prone to cell death and therefore contributed to the decrease in median cell population area observed at later time points and higher ISO doses (**Figure5E** and **H**).

As expected from the single-cell ISO concentration hypertrophy responses at 48 h, we saw a trend towards an increased proportion of grower cells with ISO stimulation compared to DMSO. In addition, the largest proportion of grower cells was seen with 1 µM ISO, which corresponds to the dose at which we saw the greatest population-level hypertrophy response (**Figures 5C** and **4C**, **D**). The relative decrease in the proportion of grower cells as ISO concentration is increased beyond 10 µM ISO also correlated with the plateauing of the population-level hypertrophy ISO concentration response (**Figure 4C** and **D**).

We hypothesized that growth and death responses may be discordant at the single-cell level, with apoptosis primarily occurring in “shrinker” cells. We quantified the proportions of cells from the three growth subpopulations that contribute to the ultrasensitive response of A5 binding with increasing ISO concentration (**Figure 5D**). At 100 µM ISO, shrinkers and stunned cells were indeed statistically more likely than growers to become A5 positive, leading us to favor a “grow-or-die” model over the “grow-and-die” model (**Figure 5E**). We consolidated these data into state-transition diagrams, which demonstrate that low ISO biases individual cells to grow, while high ISO biases cells to die. (**Figure 5F**). This concentration-dependent differential hypertrophy-apoptosis response to ISO is shown in **Supplemental Movies 4 and 5.**

Having found that apoptotic cells also tended to become smaller as “shrinkers” (**Figure 4H**), we further analyzed the growth and death kinetics of the hypertrophic subpopulations to determine additional causes for the biphasic hypertrophy concentration response observed at the population level. Overall, cells that became A5 positive exhibited a median time lag from maximal cell area to becoming A5+/PI- of 15 hours (**Figure 5G, Supplemental Figure 4**). We next asked whether the growth-death kinetics differed by hypertrophic subpopulation. Indeed, we found that stunned cells grew to their maximal extent before 24 hours while growers continued growing after 24 hours (**Figure 5H**).

The three hypertrophic subpopulations further differed in their rate of apoptosis. The grower and stunned subpopulations became A5+/PI- 11 and 13 hours after reaching their maximal cell area, respectively. Despite reaching maximal cell area early, the shrinker subpopulation persisted longer with a median time lag of 18 hours before binding A5 (**Figure 5I**). From this analysis, we conclude that the population-level biphasic hypertrophy in response to increasing ISO concentration (**Figure 4D**) arose from multiple distinct subpopulations. The shrinker subpopulation decreased in cell area and then underwent delayed apoptosis. In contrast, the grower subpopulation increased in cell area with a reduced incidence of apoptosis. These distinct subpopulation behaviors may represent an incoherent feedforward network motif between ꞵ-adrenergic receptor stimulation, hypertrophy, and apoptosis responsible for a single-cell “grow-or-die” decision.

As a validation of this “grow-or-die” conceptual model, we hypothesized that inhibiting the apoptosis pathway may redirect cell decisions to the hypertrophy pathway. Under conditions of 100 μM ISO where we had found a mixed hypertrophy-apoptosis response, treatment with caspase-3 inhibitor Z-DEVD-FMK [27] decreased cardiomyocyte apoptosis and the proportion of shrinker cells while increasing the proportion of grower cells (**Figure 5F, Supplemental Figure 5**).

We next asked whether, like ISO, PE-induced hypertrophy would also be protective from apoptosis. As shown in **Supplemental Figure 6**, PE-treated growers were less likely to die than shrinkers or stunned cells (**C**), and PE treatment increases the proportion of the grower subpopulation compared to DMSO control (**D**). Thus, the “grow-or-die” conceptual model best explains single-cell decisions in response to both ISO and PE stimulation.

### Initial cell morphology and nucleus predicts single-cell decisions of growth or death

Given the considerable single-cell heterogeneity in response to ISO stimulation, we asked whether pre-existing features of cardiomyocytes would predict their subsequent growth or death. We trained a multinomial log-linear regression machine learning model (see **Methods**) to identify cellular features that may predict single-cell decisions of growth and/or death (**Figure 6A, Supplemental Table 1**). The response variable was one of 6 pairwise combinations of the 3 hypertrophic trajectory classifications and the binary survival/death outcome. The predictor variables were the 6 cellular and 6 nuclear size, shape, and fluorescence intensity features identified by our assay as potentially relevant to cardiomyocyte apoptosis or hypertrophy (**Figures 2** and **3**). 5-fold cross-validation repeated 100 times showed that the machine learning model predicts the single-cell hypertrophy-apoptosis response to 100 µM ISO with an accuracy of 32.2±0.1% (mean±SEM) and R^2^_McFadden_ of 0.075. This accuracy was similar to that of a “random forest” model (accuracy of 32.7±0.1%, mean±SEM, **Supplemental Figure 7B**). Both multinomial regression and random forest machine learning models performed significantly better than the multinomial null model (accuracy of 20.8±0.01, mean±SEM). The similarity of accuracy in both log-linear and random forest approaches shows that differences in model structure were less important than unmeasured initial features or sources of stochasticity that cause accuracy much less than 100%.

**Figure 6.**
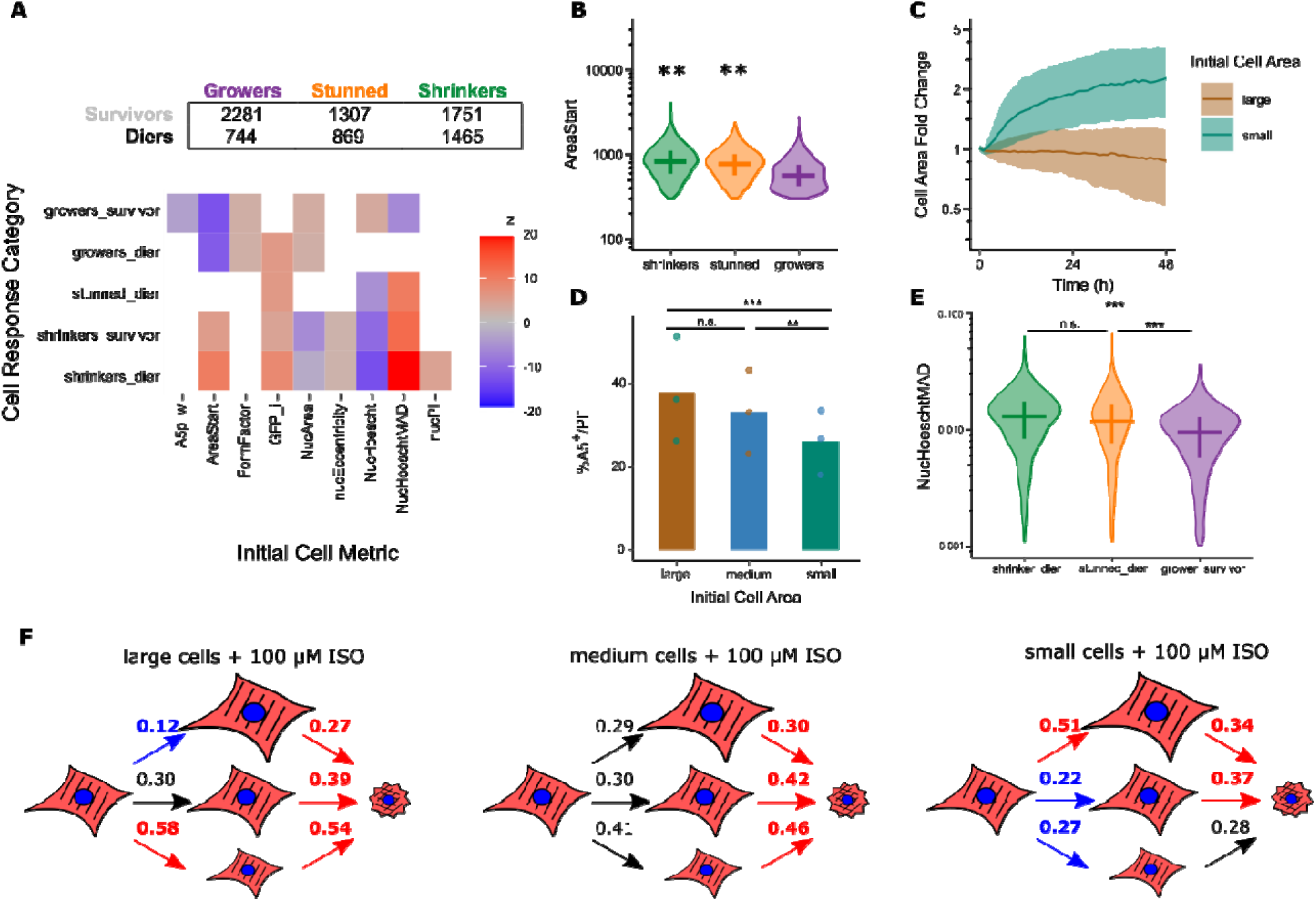
Multinomial log-linear model reveals initial cell features that predict single-cell hypertrophy-apoptosis decisions in cardiomyocytes treated with 100 μM ISO. All data is from experiments reported in Figure 4. (**A**) Significant multinomial log-linear model coefficient z-scores. The contingency table contains the number of cells in each hypertrophy-death category. The model self-validates with an accuracy of 33.26% (R^2^_McFadden_ = 0.075), with a 5-fold cross-validation accuracy of 32.33+/- 0.01% (n = 100 permutations) (**Supplemental** Figure 7). (**B**) Initial cell areas of each hypertrophic trajectory group. Solid lines indicate population median, error bars indicate interquartile range. ** p ≤ 2*10-16 vs. grower population, Kruskal–Wallis H test followed by one-sided Mann–Whitney U post-hoc test with Benjamini-Hochberg multiple testing correction, n > 2500 cells per growth category. (**C**) 48h hypertrophy dynamics of cells in either the bottom or top quartile of the initial cell area. Solid lines indicate population median, shaded areas indicate interquartile range. **(D)** 48h total apopotosis (A5+/PI-) of cells in either the bottom (<25%), intermediate (25-75%) or top (>75%) quartile of the initial cell area. *** p < 1*10^-4, ** p = 0.00267, n.s. = 0.06724. Mixed-effects ANOVA followed by Tukey HSD, 1-3 wells per condition from each of n = 3 independent cell isolations. **(E)** Initial nuclear Hoechst MAD of grower-survivor group vs. all other cells. Solid lines indicate population median, error bars indicate interquartile range. *** p < 2*10^-16, ** p = 0.0031, mixed-effects ANOVA followed by Tukey HSD, n > 1900 cells per category. **(F)** Hypertrophy and apoptosis transition state diagrams for different cardiomyocyte subpopulations. Arrow colors are relative to the DMSO diagram in Figure 5F, with red or blue indicating an increase or decrease in transition probability of > 0.05.

An advantage of the log-linear modeling approach is that it provides interpretability of the coefficient values of each classifier. The log-linear model identified several initial cell metrics that were predictive of hypertrophy, apoptosis, or both. One such metric was the initial cell area - the model predicted that initially larger cells had a higher likelihood of being stunned or shrinking and a lower likelihood of growing (**Figure 6A**). The relationships identified in the model between initial cell features and growth-death responses motivated further verification. Indeed, growers had an initial cell size that was smaller than shrinker or stunned subpopulations (**Figure 6B**). Analysis of the growth trajectory of cells in either the bottom or top quartiles of initial cell area distributions also showed that small cells had a hypertrophic growth response while larger cells did not hypertrophy significantly (**Figure 6C**). While the initial area regression coefficient did not specifically correlate with the survival status of a cell, it had an especially strong statistical association with the shrinker/dier population. Indeed, cells in the largest quartile of initial cell areas were more likely to die than those in the smallest quartile (**Figure 6D**). A parallel random forest regression model showed that the initial area was the most influential metric for model accuracy (**Supplemental Figure 7C**). Cells with small initial area were also less likely to apoptose in response to PE treatment (**Supplemental Figure 6D**). These initial cell size-dependent hypertrophy and apoptosis decisions are summarized in **Figure 6F**, where large cells are more likely to shrink and die. Our data show that with both ISO and PE treatment, hypertrophic signaling confers partial protection against apoptosis. Thus, our machine learning model of single-cell predictors of growth/death decisions further supports the “grow-or-die” model over the “grow-and-die” model.

Interestingly, this analysis identified nuclear Hoechst intensity (reflecting DNA content or nuclear permeability) and its median absolute deviation (reflecting chromatin condensation) as two nuclear metrics predictive of both cell shrinking and dying (**Figure 6A**). The median absolute deviation metric was also implicated as the second most influential metric for model accuracy by the random forest model (**Supplemental Figure 7C**). Similarly, our single-cell chromatin condensation as measured by Hoechst MAD found less condensation in grower-survivor cellscompared to other subpopulations (**Figure 6E**), which we also observed in our PE- and 56 µM ISO-treated cells at the population level (**Figure 3I** and **Supplemental Figure 8**).

## Discussion

Single-cell, multiplexed, or longitudinal data each have the potential for elucidating new causal relationships or differential cell responses to stimuli [61], [62], [63]. Most recently, single-cell and single-nucleus RNA sequencing (scRNA-seq and snRNA-seq) have identified new cardiac cell subpopulations and single-cell specific mechanisms that contribute to heart failure [24], [64], [65]. We still do not fully understand the role of functional diversity among cardiac cell types in disease progression, but recent advances in sequencing have illustrated importance in assessing cellular heterogeneity in both the healthy and failing heart.

Expression of the fetal gene program is a common surrogate for cardiomyocyte hypertrophy measurements [66], [67], but bulk measurements capture neither the changes in cell morphology nor heterogeneity of single-cell behaviors. Fixed cell cytometry approaches may capture phenotypic endpoints but lack information about the initial system state or the phenotypic trajectories necessary to elucidate decision-making processes without the use of additional system perturbations [27], [63], [68]. Previous studies of single-cell decision-making have largely been in non-myocytes and have examined decision-making for a single phenotype [28], [69], [70]. Here, our high-content microscopy platform extends previous work on automated live imaging of single-cell hypertrophy to address the challenges of measuring single-cell hypertrophy and apoptosis simultaneously. This study leveraged simple genetically encoded and biochemical fluorescent reporters to track morphological and biochemical dynamics in single cells necessary to investigate fundamental cell decisions of growth and death.

After validating our cardiomyocyte assay for hypertrophy and apoptosis measurements separately and identifying new features of those processes, we used our assay to examine the dynamics of decision-making between hypertrophy and apoptosis in response to catecholamines. Theoretical mechanisms underlying the simultaneous regulation of analog and digital phenotypic decision-making by a single signaling network have been proposed. To our knowledge, however, the relative timing of early analog hypertrophy responses and late digital apoptosis decisions represents the first experimental support of a single signaling network regulating a mixture of analog and digital phenotypic decisions [71]. Leveraging these heterogeneity dynamics in our cell populations, we identified a biphasic hypertrophic ISO concentration response at the cell-population level, implying at least two underlying, competing processes. We therefore hypothesized that different subpopulations of cells contribute to the overall population response.

Our data with both ISO and PE treatment best supports a “grow or die” conceptual model in which hypertrophic signaling is moderately protective against apoptosis. The transition-state diagrams show that while all three subpopulations undergo apoptosis at high-dose ISO treatment, the grower subpopulation is more resistant to undergo apoptosis compared to the shrinker subpopulation. Likewise, the PE-treated grower subpopulation had a decreased incidence of apoptosis. However, at low-dose ISO treatment, there is an increase in the grower subpopulation and a decrease in the shrinker subpopulation without any increase in apoptosis between those populations. Further, inhibiting caspase 3 activity re-routed cardiomyocytes from apoptosis to hypertrophy. Together, these observations of differential apoptosis among hypertrophic subpopulations undermine the “grow-and-die” model and in favor of the “grow-or-die” model under both ISO and PE treatment.

We used cyclic data-driven hypothesis testing and modeling to identify characteristic features of these different subpopulations. Our multinomial log-linear regression models sought to identify nongenetic heterogeneities in initial cell size, morphology, and fluorescence metrics at a single ISO dose that would predict dynamic, single-cell decision-making. This modeling approach had an accuracy rate similar to the more agnostic, black-box “random forest” model. This shows that differences in model structure (such as the presence of non-linear interactions between predictors) were less important than other, non-characterized factors or biochemical sources of stochasticity [72], [73]. Even so, we identified and validated a smaller initial cell size as predictive of greater hypertrophy and survival. Whether initial cell area affects in vivo cardiomyocyte hypertrophy requires additional study, but it is consistent with limited hypertrophy of aged animals subjected to stress. Surprisingly, we identified two nuclear metrics, DNA intensity (integrated Hoeschst), and chromatin heterogeneity (Hoechst MAD), that predicted cell hypertrophy and survival, which implies a potential role for transcriptional regulation of differential cell decision-making between growth and death [74], [75]. These three metrics were also predictive of hypertrophy/apoptosis decision-making with other treatments, namely PE and other ISO concentrations. The caspase inhibitor Z-DEVD-FMK decreased the “shrinker” subpopulation and increased the “grower” subpopulation, indicating that decisions toward death can be prevented.

We envision multiple avenues for extending the approach presented herein. We focused on neonatal cardiomyocytes for their feasibility of 48 hr tracking, de-differentiation/death of adult cardiomyocytes, and because substantial literature supports overall similarities with adult cells in hypertrophy-apoptosis [33], [53], [54], [58], [76]. However, there are also well-known differences with maturation, in which adult cells may be more predictive on in vivo decision-making. Tracking *in vitro* co-culture models of cardiomyocytes and fibroblasts and the relative local density of those cells would provide new insight into paracrine signaling interactions between cardiomyocytes and fibroblasts to yield new potential therapeutic targets [77]. Additional phenotypes such as cardiomyocyte proliferation could also multiplexed with our assay to examine additional single-cell decisions [48]. Our fluorescent reporter could be replaced with reporters of signaling pathway activity to provide mechanistic insight into cellular decision-making [78]. Can these decisions be tracked in vivo? Single-cell RNA-sequencing can identify sub-populations but not resolve the dynamic decision making. Light-sheet microscopy of live zebrafish has enabled testing of hypotheses previously limited to cell culture [79], [80], although these approaches have not typically had fully automated cell analysis and are less relevant to hypertrophy-apoptosis in mammalian hearts. Alternatively, the current study may motivate sophisticated dual-color lineage tracing [81] to test the “grow-or-die” conceptual model in vivo. Overall, the high-content live-cell microscopy approach presented here provides an improved understanding of cellular multi-tasking in response to stress.

## Acknowledgements

We thank members of the Saucerman lab, including B. Wissmann for technical support with cardiomyocyte isolations, P. Tan for support with CellProfiler, and L. Woo for discussions and providing feedback on this work. We also thank Chris Smolko and Kevin Janes for contributing to the eGFP lentivirus. We acknowledge Research Computing at The University of Virginia for providing computational resources and technical support.

## Author contributions

Conceptualization, B.C. and J.J.S.; Methodology, B.C., L.N. and J.J.S.; Investigation, B.C., L.N. and J.J.S.; Writing, B.C., L.N., and J.J.S.; Funding Acquisition, J.J.S.; Supervision, J.J.S..

## Methods

### Chemical Reagents

- DMSO (R&D Systems, Cat#3176)
- Staurosporine (R&D Systems, Cat#1285)
- (R)-(-)-Phenylephrine hydrochloride (R&D Systems, Cat#2838)
- Isoproterenol hydrochloride (R&D systems, Cat#1747)
- Z-DEVD-FMK (R&D Systems, Cat#2166)

### Lentivirus

Lentiviruses were prepared in 293T/17 cells (ATCC) by triple transfection with psPAX2, pMD.2G, and PLX302 eGFP as previously described [82], [83]. The plasmids were a gift from K. Janes (University of Virginia).

### Cell Culture

Cardiac myocytes were harvested from 1–2 day-old Sprague-Dawley rats using the Neomyts isolation kit (Cellutron, Baltimore, MD). All procedures were performed in accordance with the Guide for the Care and Use of Laboratory Animals published by the U.S. National Institutes of Health and approved by the University of Virginia Institutional Animal Care and Use Committee. Myocytes were cultured in plating media (Dulbecco’s modified Eagle media supplemented with 17% M199, 10% horse serum (Gibco, 16050122), 5% fetal bovine serum (HyClone FBS Thermo Scientific Cat: SH30396.03 Lot: AUD34652), 50 U/mL penicillin, 50 μg/mL streptomycin (Gibco, 15140122), 4.8mM L-glutamine(Gibco, 25030081) at a density of 20,000 cells per well of a 96-well CellBIND plate (Corning) coated with SureCoat (Cellutron). Cells were transduced with 100μL of eGFP lentivirus and plating media supplemented with 8μg/mL polybrene (Santa Cruz Biotechnology, Cat#sc-134220) at the time of plating and after 24 hours. Cells were contractile at the time of transduction. 36 hours after plating, the medium was replaced with low-serum media (phenol-red free Dulbecco’s modified Eagle media supplemented with 17% M199, 0.1% FBS, 1% ITSS, 20mM HEPES, 50 U/mL penicillin, 50 μg/mL streptomycin, 4.8 mM L-glutamine) for 24 hours of serum starvation prior to treatment. These conditions were optimized to maintain cell survival and stabilize cell area and GFP expression under DMSO treatment (see Figure 3B, Figure 3C).

### Dual hypertrophy/apoptosis assay

Cells were pre-stained and pre-treated with relevant small molecule inhibitors in low-serum media supplemented with 27 nM Hoechst 33342, 0.5 nM Propidium iodide, and 0.5× Alexa Fluor 647–conjugated A5 (Invitrogen) for 2 hours. Cells were imaged on the Operetta CLS high-content imaging system (Perkin-Elmer) using a 10× 0.3NA objective in widefield mode, which integrates over the entire cell thickness. Temperature and CO2 control options were set to 37°C and 5% CO2, respectively. Images were captured every 1 hour for 48 hours. 4 fields of view were acquired for each well.

After live-cell imaging, the cells were fixed with 4% PFA and stained with DAPI and monoclonal anti-α-Actinin antibody (Sigma, Cat#A7811). The same fields of view acquired during live imaging were again imaged.

### Image Processing

All images were processed using Matlab 2018a (MathWorks), ilastik[78], and CellProfiler[79]. Images were flat-field corrected in Matlab using reference dark field and flat-field images taken of fluorescence reference slides (Ted Pella). Corrected fluorescence signals from GFP, A5, and α-Actinin images were converted to probability maps using the pixel classification tool in ilastik [84].

An image analysis pipeline was developed in CellProfiler to automatically segment, measure, and classify the cells from the corrected images and signal probability maps. Briefly, all live timepoints and fixed immunofluorescence images were aligned to the final timepoint images. Hoechst- and propidium iodide-positive nuclei were independently segmented using Otsu thresholding. Binucleates were identified by merging touching nuclei and then shrunken by a radius of 1 pixel. GFP positive cells were identified using a watershed segmentation algorithm on the GFP probability map using the merged nuclei as starting seeds. These GFP positive cells were tracked and measured for morphological and fluorescence intensity characteristics including area, GFP intensity, A5 intensity and probability, final α-Actinin intensity and probability. Nuclei were also measured for morphological and propidium iodide and Hoechst intensities. Cells that were PI+ at the beginning of the experiment were excluded.

### Statistical Analysis

All statistical analyses were done in R version 3.6.3. The following packages were additionally used for general data preparation and visualization: tidyverse [85], multcomp [86], nlme [87], nnet [88], perturb [89], PMCMRplus [90], randomForest [91], stargazer [92] and tableplot [93]. Comparisons of single cell level metrics were performed as n = cells. Comparisons of population-level statistics were performed as n = independent cell isolations, generally with a mixed effects ANOVA with appropriate post-hoc tests. In situations in which the ANOVA was not significant but there was a predefined hypothesis, post-hoc analysis was still conducted as recommended [94]. Mixed effects ANOVA with independent cell isolations were modeled as a random effect.

### Modeling

Data were normalized prior to statistical modeling. For multinomial log-linear modelling, predictor variables were filtered for multicollinearity on the basis of their variance inflation factor (VIF) scores - removal of the raw major axis length metric reduced all VIF scores below a maximum of 2.4. We also verified the remaining collinearities with a tableplot and that those variables described sufficiently different metrics (**Supplemental Figure 7A**). For random forest modelling, default values from the randomForest package were used with the exception of 1000 trees [91].

## Supplemental Figures

**Supplemental Figure 1.**
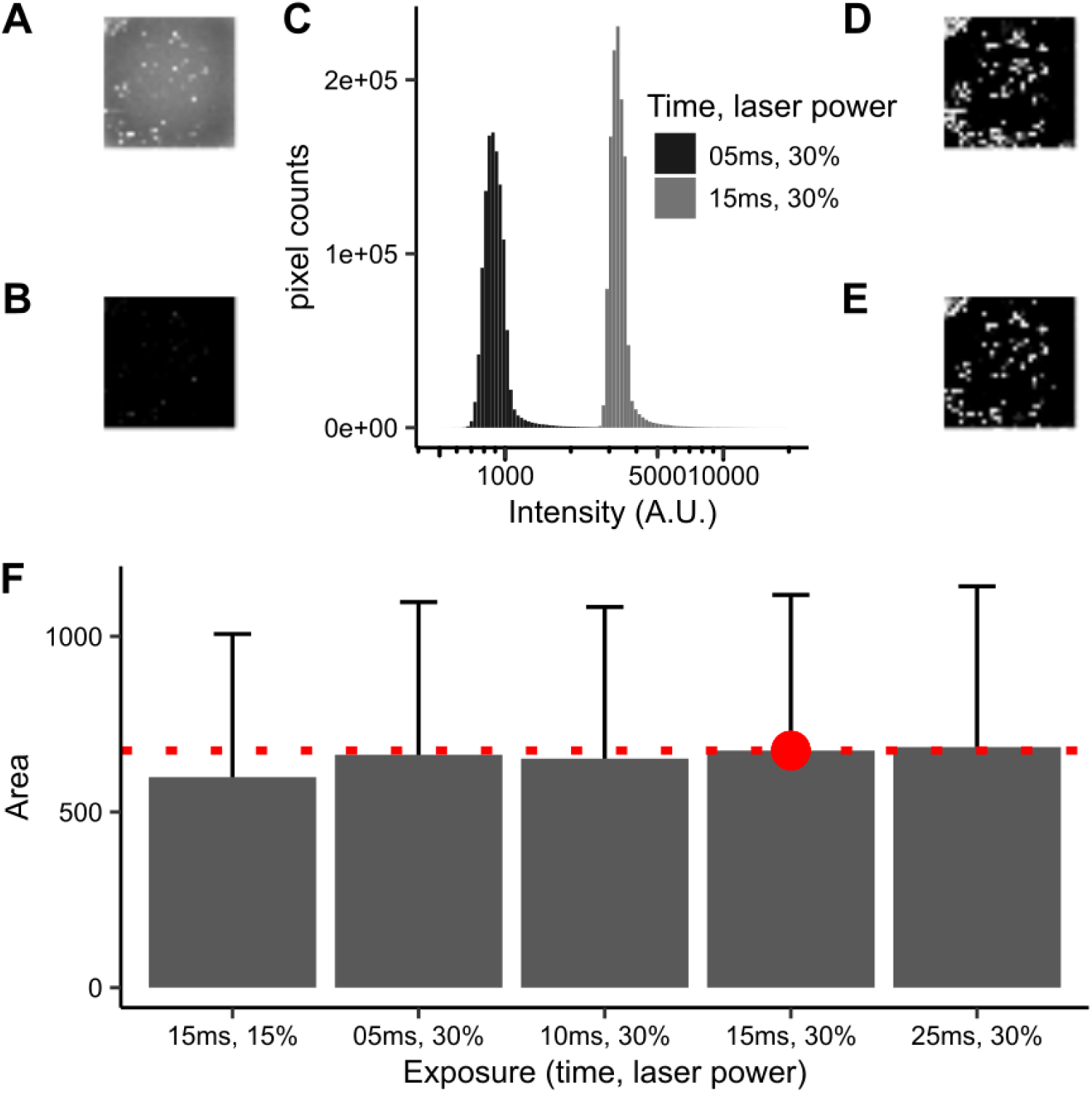
Segmentation on eGFP fluorescence probability maps is robust to 5-fold variation in signal-to-noise ratio. The same field of view of cells was taken with either 15 ms (A) or 5 ms (B) of exposure with 30% laser power. (C) Image intensity histogram of the two images. The same Ilastik random forest model was used to generate GFP fluorescence probability maps of the images taken with either 15 ms (D) or 5ms (E) of exposure. (F) Single-cell area distributions were compared with varying signal-to-noise ratios generated by varying exposure duration or laser power.

**Supplemental Figure 2.**
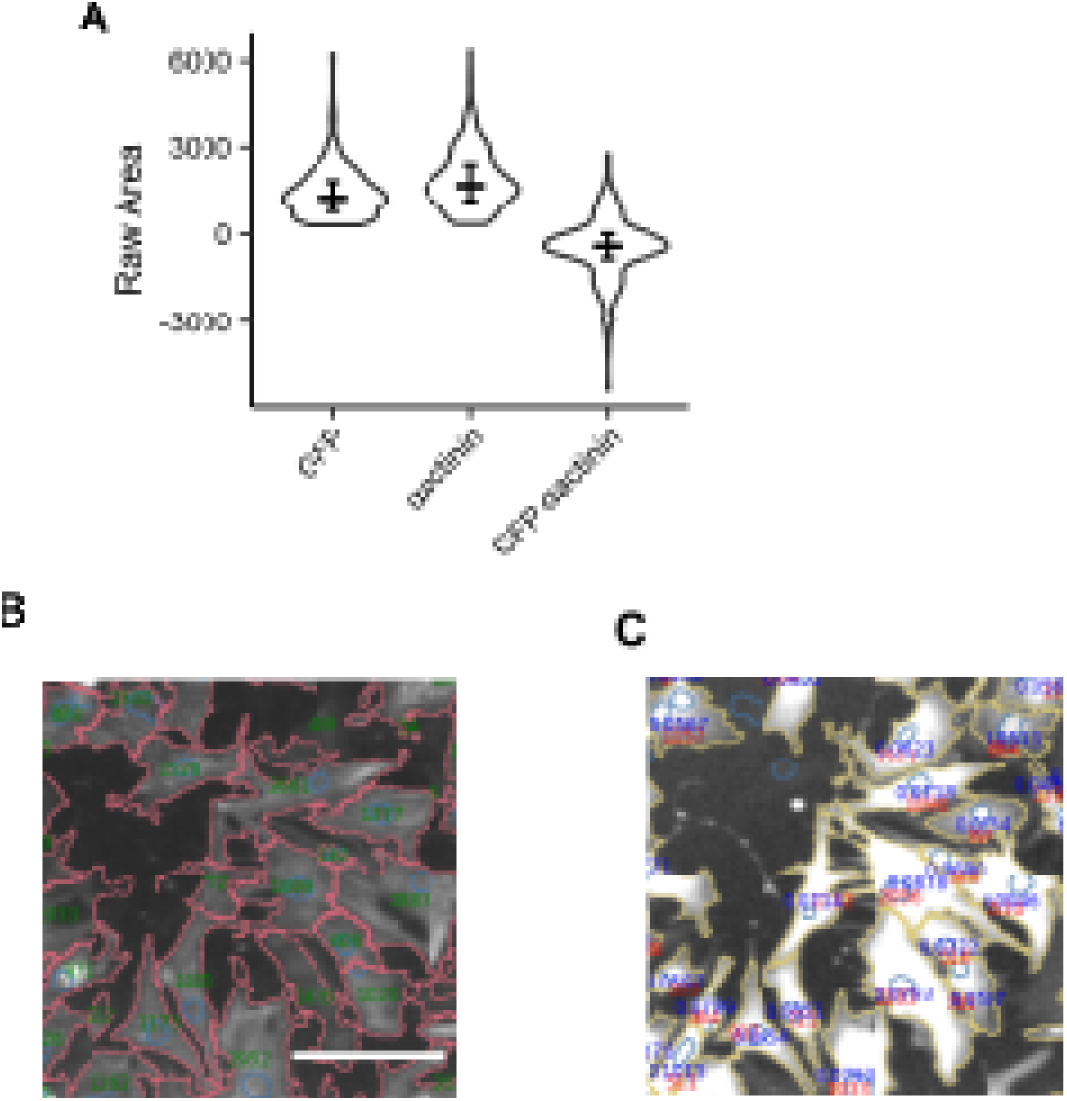
GFP watershed algorithm provides better fidelity to actual cell areas (**A**) Areas of GFP and ɑ-actinin double-positive cells segmented in GFP or ɑ-actinin channels, and the difference between the two. Solid lines indicate population median, error bars indicate interquartile range. Segmentation borders of ɑ-actinin (**B**, red), or GFP (**C**, yellow) fluorescence. Green numbers in (**B**) or red numbers in (**C**) indicate cell areas. Blue numbers in (**C**) indicate GFP fluorescence intensity.

**Supplemental Figure 3.**
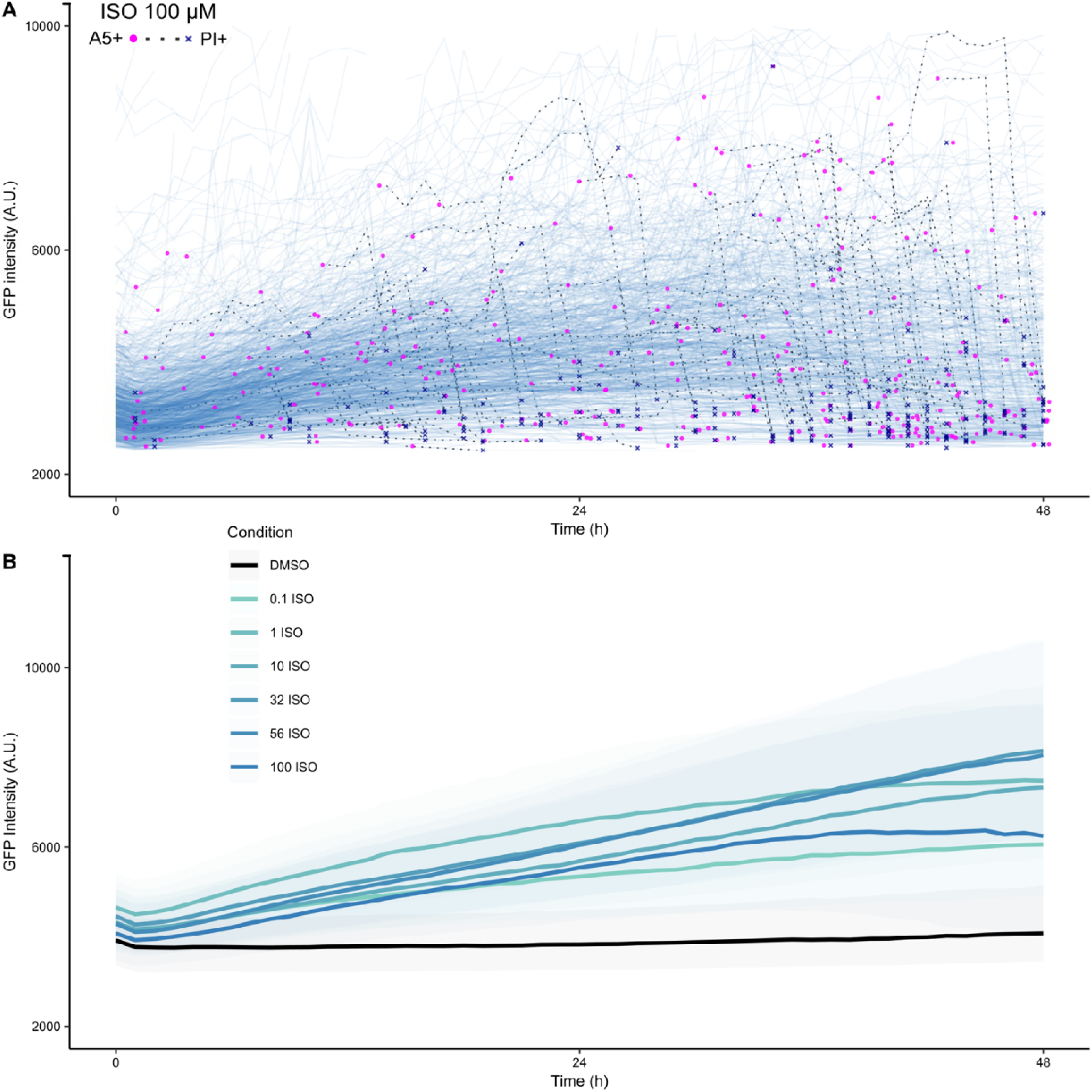
Isoproterenol induces distinct concentration response of GFP intensity, with substantial single-cell variability. (**A**) Representative single-cardiomyocyte dynamic traces of GFP intensity (representing protein synthesis) of cells treated with 100 μM ISO. Magenta dots indicate onset of A5+/PI- binding, with the blue ‘X’ marker indicating eventual PI binding in that same cell. (**B**) eGFP dynamics of cells treated with various ISO concentrations vs. DMSO vehicle control. Solid lines indicate population median, shaded areas indicate interquartile range.

**Supplemental Figure 4.**
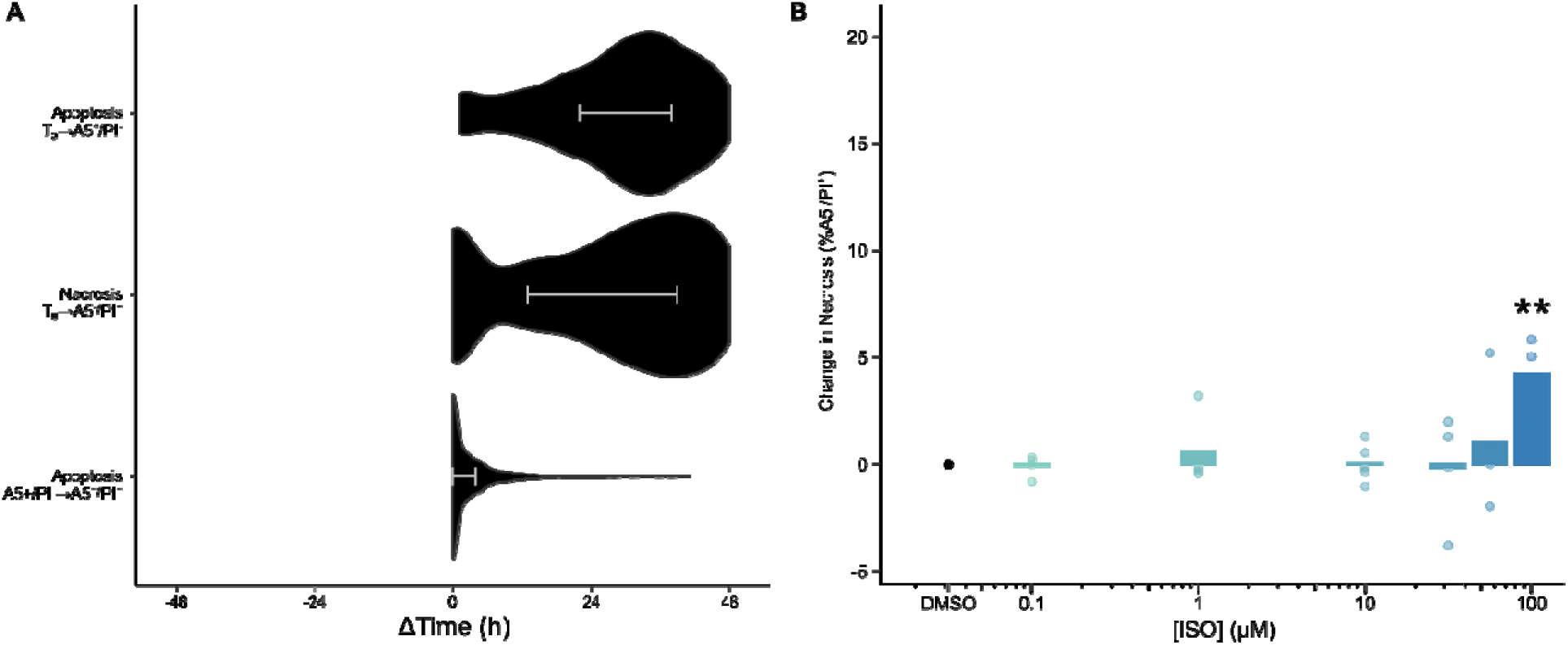
Single-cell hypertrophy-necrosis tracking in response to isoproterenol. (**A**) Single-cell time lag from maximal cell area to apoptosis or necrosis, or from A5+/PI- to A5+/PI+ for 100 μM ISO. (**B**) ISO concentration of necrosis (A5-/PI+) through 48 h. * p < .05, *** p < .001 vs DMSO control; mixed effects ANOVA followed by Dunnett post hoc test, same wells as in **Figure 4C** with n = 3-5 independent cell isolations.

**Supplemental Figure 5.**
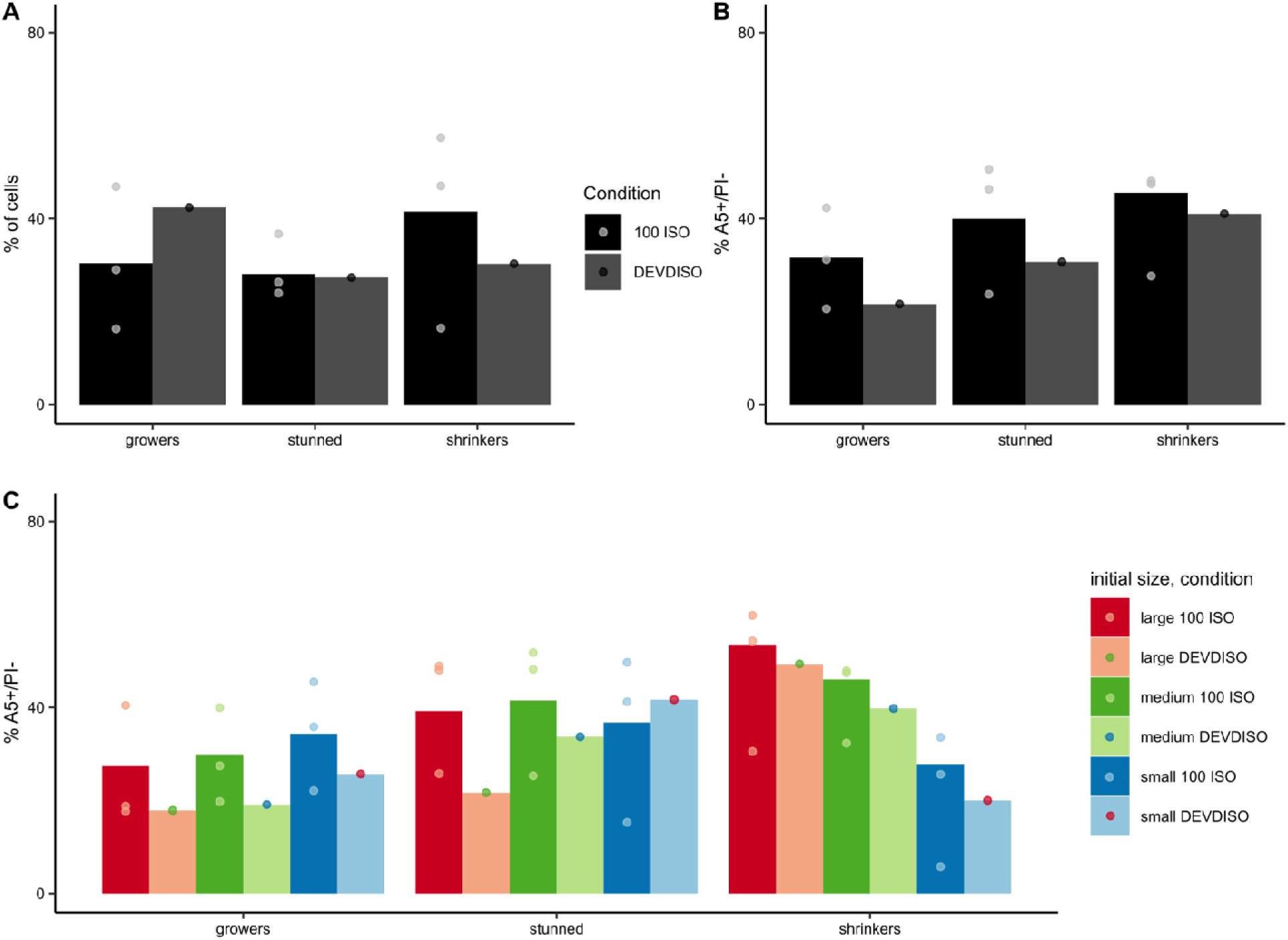
Caspase-3 inhibitor Z-DEVD-FMK inhibits ISO-induced apoptosis and increases grower cell subpopulation. (**A**) Addition of 20 µM Z-DEVD-FMK increases the proportion of grower cells while decreasing the proportion of shrinker cells. (**B**) Addition of Z-DEVD-FMK decreases the death rates of each growth subpopulation. (**C**) Addition of Z-DEVD-FMK decreases the death rates of each growth subpopulation except for the initially small, stunned growth subpopulation.

**Supplemental Figure 6.**
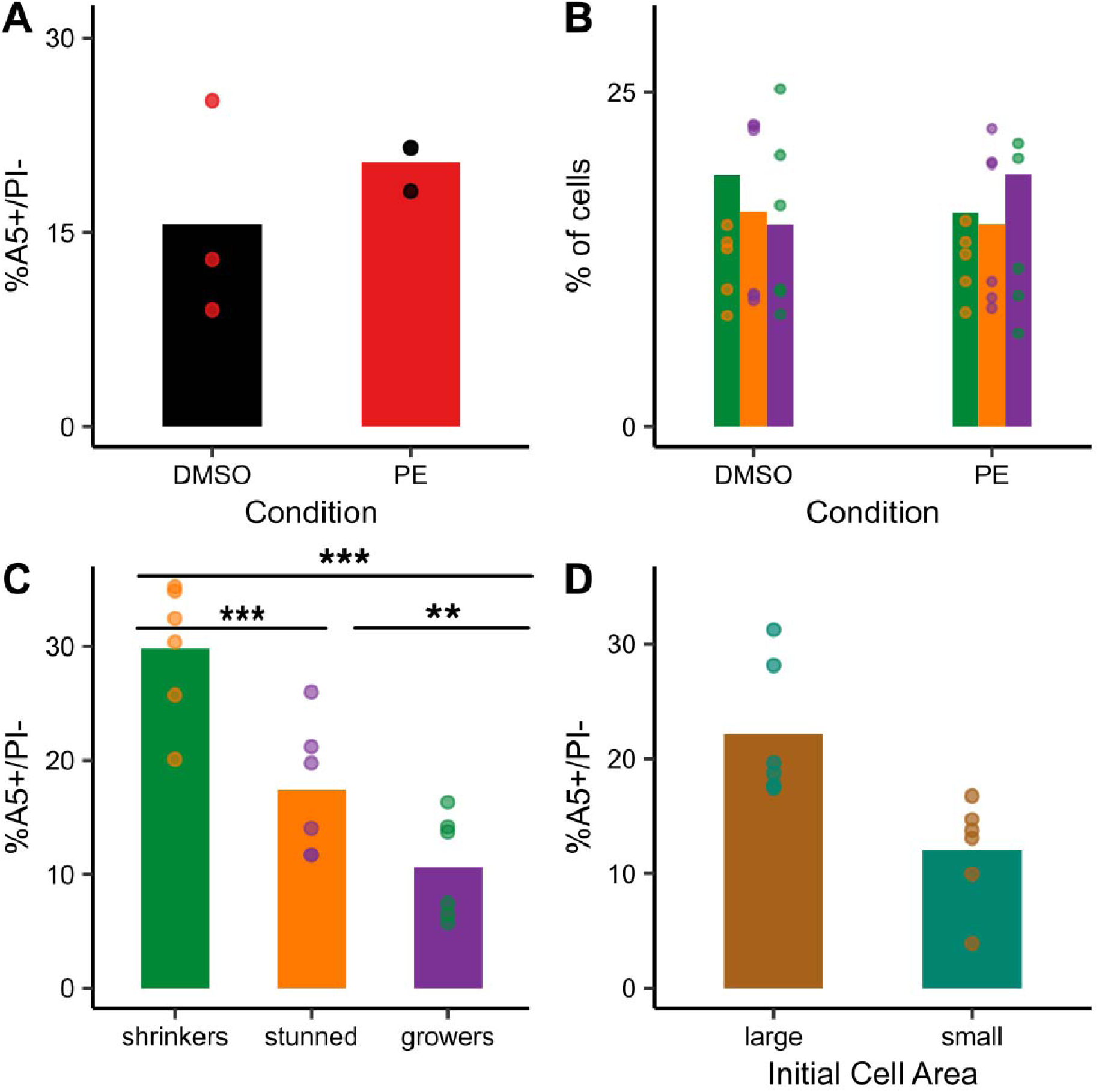
Phenylephrine increases apoptosis-resistant grower cell subpopulation. (**A**) Addition of 50 μM PE does not affect rate of cell death. p = 0.427, one-sided Welch’s paired *t*-test, 1-3 wells for each condition from n = 3 independent cell isolations. (**B**) Addition of PE increases the proportion of grower subpopulation. p = 0.121, mixed effects ANOVA with post-hoc Dunnett’s test and Bonferroni correction. 1-3 wells for each subpopulation per each condition from n = 3-4 independent cell isolations. (**C**) While the overall mixed-effects ANOVA was not statistically significant (p=0.063) the greater-powered Tukey HSD test showed that the grower subpopulation was more resistant to apoptosis than the shrinker subpopulation (*** p < 0.001, ** p = 0.00541, 1-3 wells for each subpopulation from n = 3 independent cell isolations). (**D**) PE-treated cells in the largest quartile of initial cell areas were more likely to die than cells in the smallest quartile. n.s. p = 0.998, one-sided Welch’s paired *t*-test, 1-3 wells for each condition from n = 3 independent cell isolations.

**Supplemental Figure 7.**
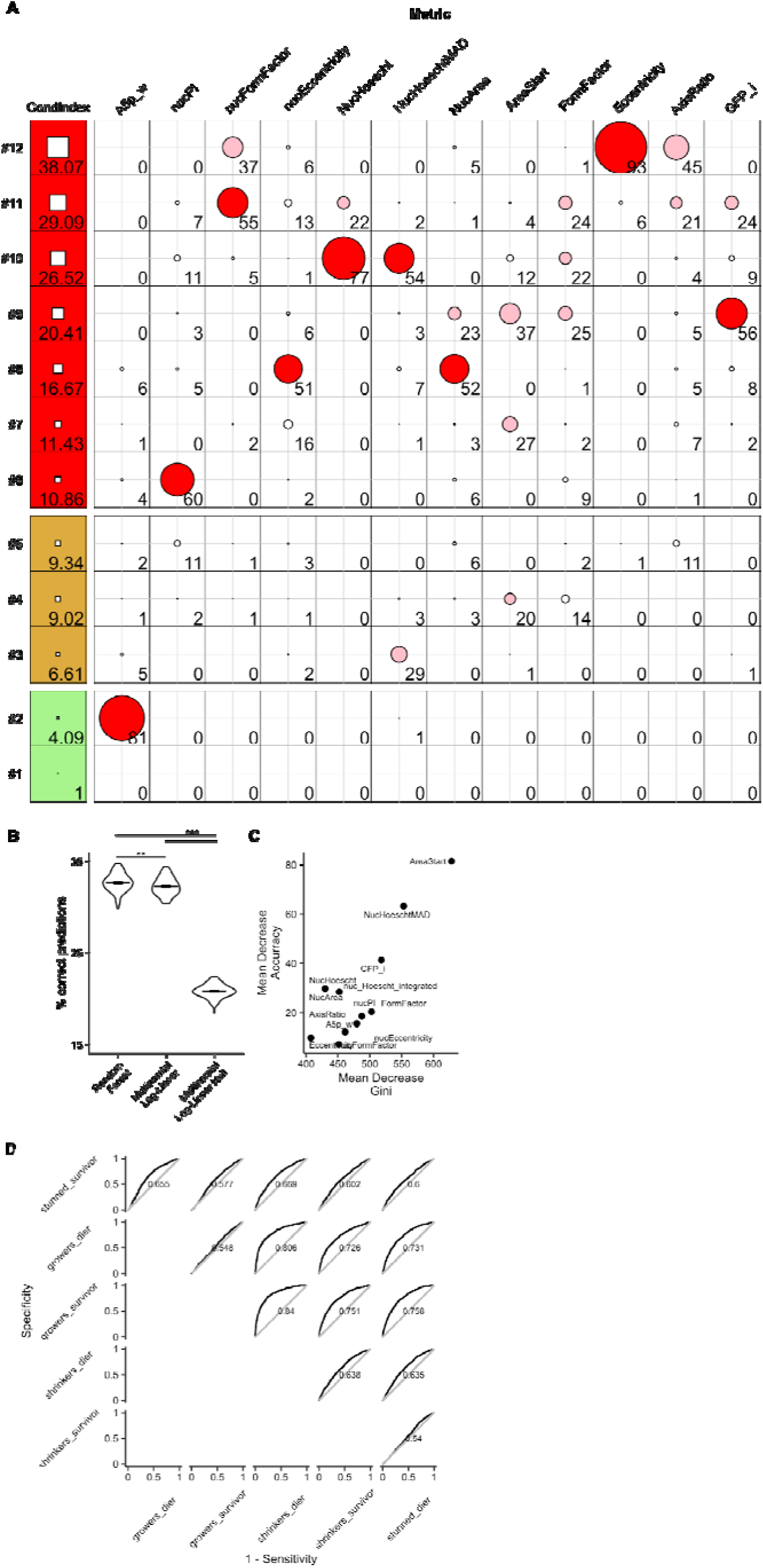
Colinearity of predictor variables and comparison of model performances. (**A**) Tableplot of Condition indices and coefficient variance proportions [90]. Briefly, condition indices represent scaled eigenvalues of the correlation matrix among the predictors variables. The condition index box is green for indexes less than 5, yellow if less than 10, and red if greater than 10. Each row represents an eigenvector with the “metric” columns corresponding to each predictor variable’s proportion of variance scaled to 100. (**B**) 5-fold cross-validation accuracy. Error bars represent mean ± SEM. **p = 0.002, ***p < 2*10^-22^, repeated-measured ANOVA followed by paired T-test with Benjamini-Hochberg correction for multiple comparisons, n=100 random partitions. (**C**) Random forest predictor variable importance measures. (**D**) Multiclass and pairwise AUC curves.

**Supplemental Figure 8.**
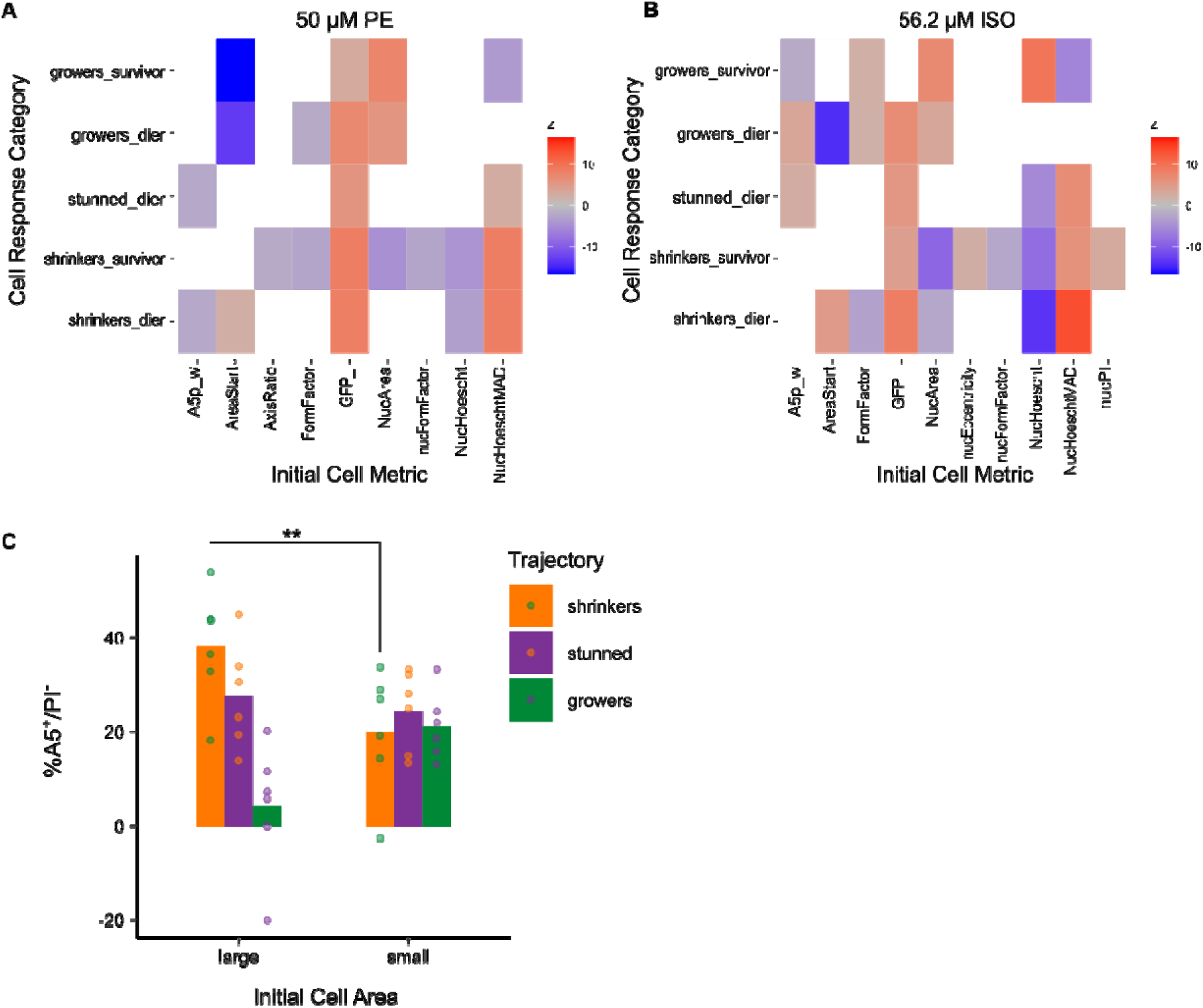
Multinomial log-linear model coefficients for PE and 56.2 μM ISO. (**A**) Model for 50 μM PE. (**B**) Model for 56.2 μM ISO. (**C**) Apoptosis rate by initial cell area and growth trajectory. ** p = 0.006268, one-sided Welch’s paired *t*-test, 1-3 wells for each condition from n = 3 independent cell isolations.

## Supplemental Tables

**Supplemental Table 1.** Multinomial log-linear regressions predicting hypertrophy-apoptosis phenotype likelihood relative to the stunned survivor phenotype. Values are represented as the regression coefficients and standard errors in parentheses. Akaike Information Criterion = 30818.4

|  | *Dependent variable:* |  |  |  |  |
| --- | --- | --- | --- | --- | --- |
|  | growers_dier | growers_survivor | shrinkers_dier | shrinkers_survivor | stunned_dier |
|  | (1) | (2) | (3) | (4) | (5) |
| A5p_w | -0.247 | -0.601*** | -0.089 | -0.108 | 0.058 |
|  | (0.200) | (0.173) | (0.178) | (0.166) | (0.194) |
| nucPI | 0.615 | 0.326 | 2.405*** | 1.310** | 0.313 |
|  | (0.648) | (0.572) | (0.552) | (0.538) | (0.636) |
| nucFormFactor | 0.566 | -0.159 | 0.433 | -0.679* | 0.553 |
|  | (0.452) | (0.372) | (0.397) | (0.365) | (0.439) |
| nucEccentricity | -0.160<br>(0.313) | -0.201<br>(0.265) | 0.471<sup>*</sup><br>(0.282) | 0.369<br>(0.265) | 0.552<sup>*</sup><br>(0.311) |
| NucHoescht | -0.162<br>(0.889) | 2.559<sup>***</sup><br>(0.762) | -11.594<sup>***</sup><br>(0.853) | -7.476<sup>***</sup><br>(0.789) | -5.250<sup>***</sup><br>(0.896) |
| NucHoeschtMA<br>D | 1.540<sup>*</sup><br>(0.786) | -4.929<sup>***</sup><br>(0.688) | 14.391<sup>***</sup><br>(0.717) | 8.033<sup>***</sup><br>(0.666) | 8.553<sup>***</sup><br>(0.763) |
| NucArea | 2.486<sup>***</sup><br>(0.725) | 2.450<sup>***</sup><br>(0.588) | -1.229<sup>*</sup><br>(0.632) | -3.251<sup>***</sup><br>(0.586) | 0.570<br>(0.700) |
| AreaStart | -9.710<sup>***</sup><br>(0.685) | -7.923<sup>***</sup><br>(0.532) | 4.959<sup>***</sup><br>(0.461) | 2.037<sup>***</sup><br>(0.452) | 0.750<br>(0.531) |
| FormFactor | 0.802<sup>***</sup><br>(0.309) | 0.685<sup>***</sup><br>(0.265) | -0.017<br>(0.275) | 0.151<br>(0.262) | 0.311<br>(0.301) |
| Eccentricity | 0.018 | 0.277 | 0.220 | -0.099 | -0.104 |
|  | (0.404) | (0.345) | (0.364) | (0.336) | (0.392) |
| AxisRatio | -1.123 | -0.701 | -0.443 | 0.357 | -0.034 |
|  | (0.792) | (0.634) | (0.629) | (0.572) | (0.694) |
| GFP_i | 4.564*** | -0.182 | 5.490*** | 2.564*** | 4.590*** |
|  | (0.718) | (0.688) | (0.640) | (0.633) | (0.697) |
| Constant | -1.182 | 1.363* | -2.813*** | 1.057 | -3.031*** |
|  | (0.903) | (0.769) | (0.802) | (0.745) | (0.882) |

\*p<0.1; \*\*p<0.05; \*\*\*p<0.01

## Supplemental Movies

**Supplemental Movie 1. Live single cell tracking of DMSO-treated neonatal rat cardiomyocytes across 48 hours.** Frames were sampled once per hour (0-48 hrs, 49 frames total). Left panel shows merged raw images of Hoechst (blue) and GFP (green) stains. Middle panel shows automated CellProfiler segmentation and nuclei tracking of cells. Right panel shows merged Annexin V (yellow) and PI (magenta) stains. Scale = 100µm.

**Supplemental Movie 2. Live single cell tracking of staurosporine (STS)-treated neonatal rat cardiomyocytes across 48 hours.** Cardiomyocytes treated with µM STS show increased rates of apoptosis and cell death.

**Supplemental Movie 3. Live single cell tracking of phenylephrine (PE)-treated neonatal rat cardiomyocytes across 48 hours.** Cardiomyocytes treated with 5o µM PE induce a hypertrophic response shown by an increase in cell area and GFP expression

**Supplemental Movie 4. Live single cell tracking of 10** µ**M isoproterenol (ISO)-treated cells across 48 hours.** Cells treated with 10µM ISO show mixed responses of apoptosis and hypertrophy.

**Supplemental Movie 5. Live single cell tracking of 100** µ**M ISO-treated cells across 48 hours.** Cardiomyocytes treated with 100 µM ISO show both hypertrophy and apoptotic responses but shift towards apoptosis at the end.

